# Bio-Inspired Ultrasound-Driven Ultrafast Soft Microgripper

**DOI:** 10.1101/2025.07.30.667446

**Authors:** Chengxi Zhong, Vincent Winderol, Khemraj Gautam Kshetri, Cornel Dillinger, Tommaso Bianchi, Zhan Shi, Song Liu, Justus Schnermann, Raphael Wittkowski, Nitesh Nama, Daniel Ahmed

## Abstract

Acoustically actuated soft matter offers potential for agile microscale manipulation, yet acoustic-soft matter interaction at the microscale remains poorly understood. Here, we explore the mechanism of ultrasound-soft matter interaction by developing a bio-inspired ultrasound-driven soft hydrogel microgripper. This exploration allows to delve deeper into the understanding of nonlinear dynamics, mode coupling, and energy transfer. The developed microgripper (≤ 120 µm) overcomes key challenges of existing grippers, including complex fabrication, reliance on additives or external wiring, rigid structures, slow or poorly controllable responses, and risks of sample damage or contamination. Interacting with acoustic actuation, soft microgrippers oscillate and deform, while adjusting acoustic parameters and microgrippers’ structures allows for programmable interactions. The optimized acoustic actuation of the soft microgripper enables precise, ultrafast (∼2 ms) gripping and handling of distinct delicate objects. This work advances the integration of soft matter with acoustic actuation especially at the microscale, offering a versatile, reliable, and scalable solution for microrobotics, targeted drug delivery, and lab-on-a-chip applications.

**Teaser:** Acoustic-soft matter interaction validated on bio-inspired ultrasound-driven ultrafast soft microgrippers.

## Introduction

Soft matter exhibits remarkable responsiveness to external stimuli (*1, 2*), enabling versatile control for microscale manipulation. For instance, magnetic fields have been widely used to actuate soft matter by embedding ferromagnetic particles (*3–6*), allowing programmable deformation under applied fields. Light fields also have been used to actuate and move soft matter, enabling fast, efficient, and controlled shape changes (*7*). Acoustic fields offer another promising method (*8–14*), providing non-contact, rapid, and biocompatible actuation without requiring embedded magnetic materials, thereby preserving the intrinsic softness and uniformity of soft matter. However, the fundamental mechanisms governing acoustic-soft matter interaction, particularly at the microscale, remain poorly understood. This knowledge gap limits the development of acoustically driven soft microscale actuators. In this work, we investigate acoustic-soft matter interaction and validate its potential by ultrasound-driven soft microgripper, demonstrating how tether-free soft microgripper achieves ultrafast and programmable control under acoustic actuation.

Microgrippers are essential tools for applications such as single-cell analysis (*10, 15, 16*), microsurgery (*17*), lab-on-a-chip systems (*14, 18*), and targeted delivery (*19*), where high precision, speed, and biocompatibility are critical. However, conventional microgrippers made from rigid materials (e.g., silicon or metal) often exhibit mechanical mismatch with soft, delicate biological targets such as cells and tissues—leading to poor contact, sample damage, or unreliable manipulation. These limitations highlight the need for soft, high-performance microgrippers that are not only precise and fast, but also compliant, gentle, and capable of operating effectively in confined or fluidic environments, as required in emerging biomedical and microrobotic applications (*20, 21*).

Various microscale gripping techniques have been developed (*22*), each employing distinct actuation principles. Rigid mechanical microgrippers utilizing MEMS structures or elastic flexures (*23–25*) provide high precision but face challenges including complex fabrication, potential sample damage from direct contact, and microscale friction or stiction issues. Pneumatic, hydraulic or vacuum-driven microgrippers (*26–29*) offer relatively rapid actuation but require intricate microchannels and micropumps that limit miniaturization and raise fluid leak risks. Electrostatic (*30, 31*) and magnetic microgrippers (*32, 33*) enable untethered operation but depend on specialized additives that may compromise biocompatibility. Optical tweezers (*34–36*) provide contactless manipulation across diverse materials, but they are constrained by limited penetration depth and potential thermal damage from high-intensity lasers. Lastly, thermally triggered microgrippers (*37–42*) can achieve significant deformations through shape-memory, swelling-induced actuation or mechanical property changes, yet their slow response times and heat generation pose risks for biological applications. Although soft matter enables conformal gripping and improved safety, current thermal-actuation approaches typically operate at larger scales (millimeter or centimeter) and face material compatibility limitations, highlighting the need for alternative actuation strategies that combine microscale precision, rapid response, and biological safety.

Hydrogels have emerged as a promising material for microscale machines (*43*), soft robotics (*44–50*), and artificial muscles (*51*) due to their biocompatibility, biodegradability, and tunable mechanical properties. Their responsiveness to diverse external stimuli, such as pH (*48, 49*), biochemistry (*52, 53*), temperature (*39, 51*), light (*46, 54*), electric fields (*55, 56*), magnetic fields (*3, 50, 57*), and acoustic fields (*43, 58*), enables dynamic and programmable shape morphing for precise manipulation. A representative approach involves magnetic actuation, where ferromagnetic domains are pre-embedded within the hydrogel using direct ink writing (DIW) technique, achieving sub-second responses but face size and resolution limitations from nozzle constraints (*3*). Alternative strategies like thermally actuated shape-memory hydrogels (> 500 µm) (*40*) combine near-infrared response with geometrical interlocking, yet remain constrained by size and speed. Acoustic stimulation has recently gained attention as a contactless, tunable activation method (*58, 59*). While ultrasound-induced heating has enabled centimeter-scale (∼ 2-3 cm) hydrogel grippers (*58*), most existing designs rely on slow (minutes-scale) swelling mechanisms in response to thermal or pH changes, which risk sample damage and offer limited control. In contrast, purely acoustic actuation bypasses these limitations, enabling rapid, reversible deformation (*43*). Despite these advantages, significant gaps remain: (i) purely acoustic hydrogel microgrippers are scarce, (ii) the fundamental acoustic-hydrogel interaction at microscales is poorly understood, and (iii) current microgripper designs lack the combination of response speed, control precision, programmability, mass production, scalability, and biocompatibility needed for practical applications.

Inspired by the natural Venus flytrap and *vorticella*, this work introduces a fundamentally new concept in tether-free, programmable, ultrafast responsive, biocompatible artificial microcilia using acoustically activated soft hydrogel structures, namely “SonoGripper”. This SonoGripper (≤ 120 µm), fabricated via single-step UV photopolymerization with biocompatible polyethylene glycol diacrylate (PEGDA) hydrogel, demonstrates remarkable responsiveness to acoustic excitations with a 2 ms reaction time. Through systematic characterization of acoustic-hydrogel interactions, we establish programmable control over gripping behaviors by tuning geometric parameters and ultrasound parameters, eliminating the need for additives or rigid components. SonoGripper enables gentle, fast, contamination-free manipulation of diverse delicate objects, making it suitable for microrobotics, targeted drug delivery, and lab-on-a-chip systems. This work not only advances the fundamental understanding of acoustic-soft matter interactions but also establishes a versatile platform for next-generation microscale manipulators.

## Results

### Bio-inspired SonoGripper fabricated by photopolymerization

Inspired by the Venus flytrap, which rapidly snaps its two hinged lobes to capture insects upon sensing external stimuli, and the *Vorticella*, which employs rapid cilia oscillation to remotely attract microorganisms (Fig. 1A), we developed “SonoGripper”, a hydrogel-made microgripper featuring two gripping arms with outward-pointed sharp edges at the arm tips (Fig. 1B left). To ensure stability during gripping, the arms are attached to a bulk body. Upon external ultrasound excitation, the SonoGripper’s arms undergo rapid oscillation and deformation, generating acoustic streaming vortices near the sharp edges (Fig. 1B right). These vortices with induced suction effects cause remote object attraction and controlled arm closure, enabling grip and encapsulate objects. The increased deformation of SonoGripper arms store elastic potential energy, thus, when the acoustic excitation is switched off, the arms swiftly revert to the original states.

**Fig. 1.**
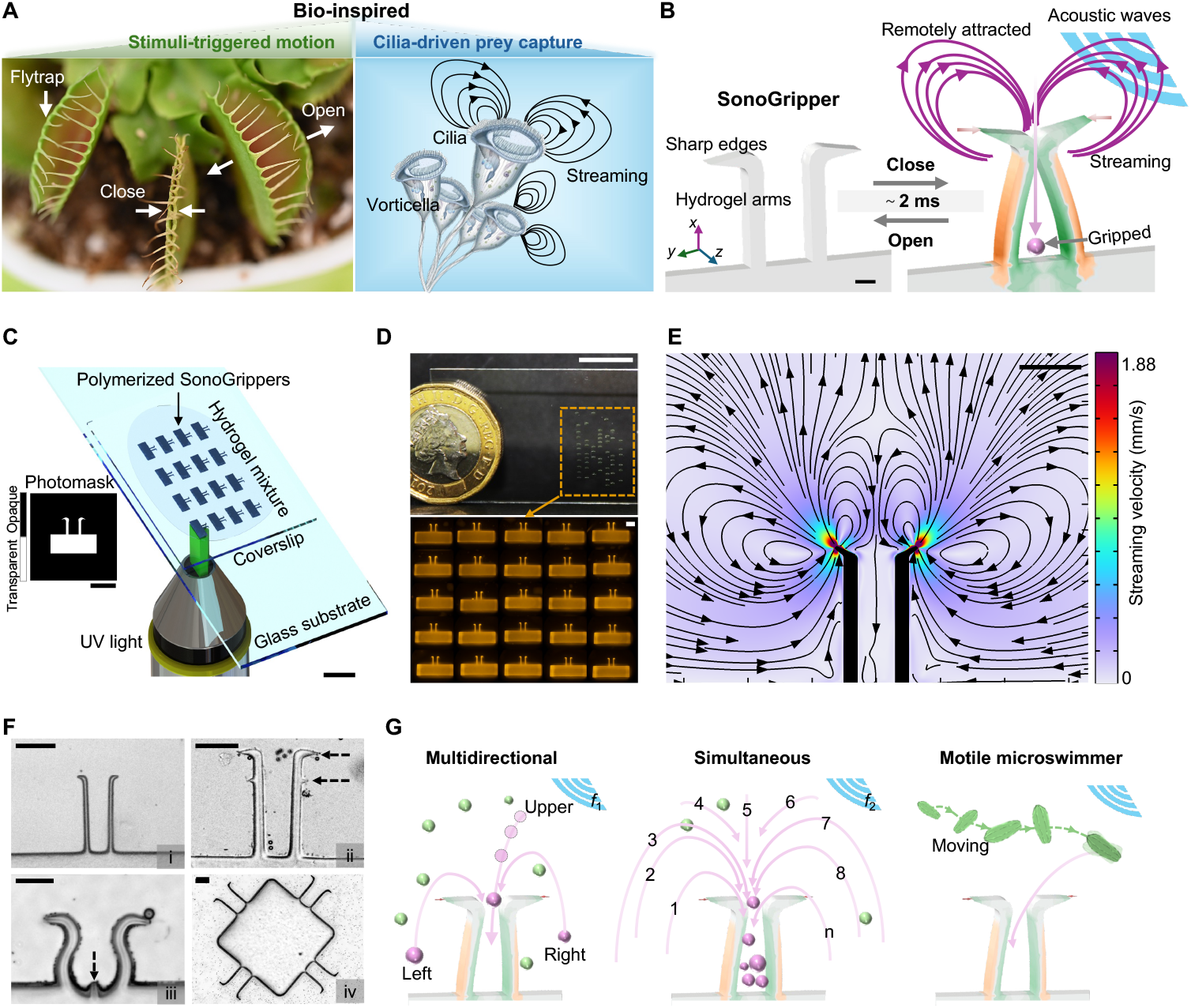
SonoGripper inspired by natural Venus flytrap (Dionaea muscipula) and *Vorticella*. (**A**) Photographs show that the flytrap rapidly snaps its two hinged lobes to capture insects upon sensing external stimuli, while the *vorticella* employs cilia’s rapid oscillation to generate fluid streaming to remotely attract microorganisms. (**B**) Schematic illustration of our developed bio-inspired SonoGripper, which mimics flytrap’s stimuli-triggered motion and *vorticella*’s cilia-driven remote attraction mechanism. Under acoustic waves, SonoGripper’s arms undergo rapid oscillation and deformation, generating acoustic streaming vortices around their outward-pointed sharp edges. These vortices facilitate object attracting, arm closure, and object encapsulation. When the acoustic excitation is switched off, the SonoGripper’s arms swiftly revert to their original states. Scale bar is 10 µm. (**C**) One-step UV-light photopolymerization developed on an inverted microscope setup (scale bar is 5 mm) utilizes UV light and a photomask (scale bar is 100 µm) to fabricate SonoGripper. The photomask with transparent and opaque regions determines local UV exposure to selectively pattern a soft geometry from photosensitive hydrogel mixture. (**D**) Image of high throughput array of SonoGrippers on a thin glass substrate (scale bar is 10 mm) and its optical microscope image (Scale bar is 50 µm). (**E**) The simulation shows a SonoGripper (100 µm in length, 7 µm in width, 20 in thickness) triggered by ultrasound, creating acoustic streaming vortices. Scale bar is 50 µm. (**F**) Optical microscope image snapshots of fabricated SonoGripper variants with different structures including (i) basic version, (ii) double-edged arm tips, (ii) curved arms with a central edge, and (iv) multi-gripper arrays combined on a bulk body. Scale bar is 50 µm. (**G**) Schematic illustration demonstrates the SonoGripper can fast, remotely, multidirectionally, and simultaneously attract and grip diverse targets with varying sizes, shapes, and mobilities.

We fabricated SonoGrippers using a custom-built ultraviolet (UV) photopolymerization technique developed on an inverted microscope setup (Fig. 1C, Materials and Methods). This technique utilizes high-resolution (20,000 dpi) photomasks designed in AutoCAD software and UV light exposure controlled by a shutter and LabView software to induce a single-step polymerization process. The photomasks (Fig. S1) contain transparent and opaque regions that determine local UV exposure, selectively patterning a soft geometry from photosensitive hydrogel mixture. The hydrogel mixture comprised 85 vol% polyethylene glycol diacrylate (PEGDA, MW 700, Sigma-Aldrich) and 15 vol% photo-initiator (2-hydroxy-2-methyl1-phenyl-propan-1-one, Sigma-Alderich). A 10 µL droplet of hydrogel mixture was placed on a glass substrate and covered with a coverslip to ensure uniform thickness and consistent polymerization. This glass substrate was then mounted on the microscope while the microscope system was carefully calibrated to precisely focus UV light on the hydrogel plane, ensuring accurate alignment and polymerization in the desired regions. To balance between structural integrity and flexibility, we estimated the Young’s modulus of SonoGrippers fabricated under different exposure times using atomic force microscopy (AFM) with the force-displacement measurements (Fig. S2, supplementary text). The results indicate that increased exposure time enhances stiffness. Insufficient exposure resulted in excessively soft structures prone to breakage under high excitation voltages, while excessive exposure produced overly thick and rigid structures, impairing their ability to oscillate effectively. An optimized UV exposure time of 40-50 milliseconds was used to fabricate SonoGrippers with dimensions of 50∼120 µm in length, 5∼20 µm in width, and 20∼25 µm in thickness. The outward-pointed sharp edges at the arm tips, measuring 15 µm long with a 30° slope, are crucial for generating acoustic microstreaming to attract and grip objects. The slender, elastic hydrogel arms enable rapid, reversible deformation upon ultrasound actuation, facilitating efficient closure and gripping. Further fabrication details are provided in the Materials and Methods.

Our high-throughput fabrication allows the production of up to 1000 SonoGrippers per batch within a minute, demonstrating its mass-producible ability (Fig. 1D). Additionally, by adjusting the photomask designs, we achieved SonoGripper variants including basic version, double-edged arm tips, curved arms with a central edge, and multi-gripper arrays combined on a single bulk body (Fig. 1F). The microstreaming under ultrasound exposure, illustrated by numerical simulations of a SonoGripper with dimensions of 100 µm in length, 7 µm in width (Fig. 1E), facilitate fast, multidirectional, and simultaneously high-throughput object attraction, where the object can feature with varying sizes, shapes, and mobilities (Fig. 1G).

### Experimental ultrasound setup

We characterized the SonoGripper using an experimental setup that includes a circular piezoelectric transducer (PZT, Murata 7BB-27-4L0) affixed to the side of a glass substrate with a two-component epoxy adhesive. The PZT was driven by a function generator (AFG3011C, Tektronix) connected to a signal amplifier (HF-Amplifier, Digitum-Elektronik), generating acoustic waves with tunable excitation voltages (1-20 V_PP_) and frequencies (5-100 kHz). We dispensed a droplet of deionized water (50-80 µL) onto the fabricated SonoGripper using a pipette and carefully placed a coverslip on top of deionized water to create a stable aqueous environment next to PZT. The setup was then mounted on an inverted optical microscope, with the SonoGripper site monitored using a high-speed camera (Chronos 1.4, Kron Technologies). To analyze controllable SonoGripper dynamics and generated acoustic microstreaming profiles under varying SonoGripper geometrical properties and ultrasound parameters, we introduced tracer particles (monodisperse polystyrene microspheres, 2–15 µm) into the aqueous environment. Streaming velocities were quantified using particle image velocimetry (PIV) software (PIVLab) (*60*), while high-speed image sequences were processed in image analysis software (ImageJ) (*61, 62*) to superimpose frames and track object movements. Additional experimental details are provided in Materials and Methods.

Deformation Closed

### Interaction between acoustic excitation and SonoGripper

We studied the dynamic interaction between SonoGripper and acoustic stimuli by analyzing gripper’s oscillation, deformation, and induced microstreaming under ultrasound exposure (Fig. 2A). Under acoustic actuation, the (first-order) oscillatory pressure on the inner and outer sides of the SonoGrippers differ, as shown in Fig. 2B. When time-averaged, these differences lead to an attractive force between the grippers that drives further elastic deformation, pulling the arms towards each other, and finally completing closure. A SonoGripper with dimensions of 100 µm in length, 10 µm in width, and 20 µm in thickness exhibited an ultrafast and repetitive oscillation (Fig. 2C, Fig. S3 and Movie S1), which is also quantified by tracking *y*-axis oscillation displacements at three sampled positions along the SonoGripper arm (50, 60, and 70 μm from the fixed end) (Fig. 2D). At 50 and 60 μm, oscillation remained periodic and relatively stable at ∼2.5 μm oscillation displacement. Near the arm tip (at 70 μm), the oscillation displacements progressively increased, eventually causing arms contact and SonoGripper closure.

**Fig. 2.**
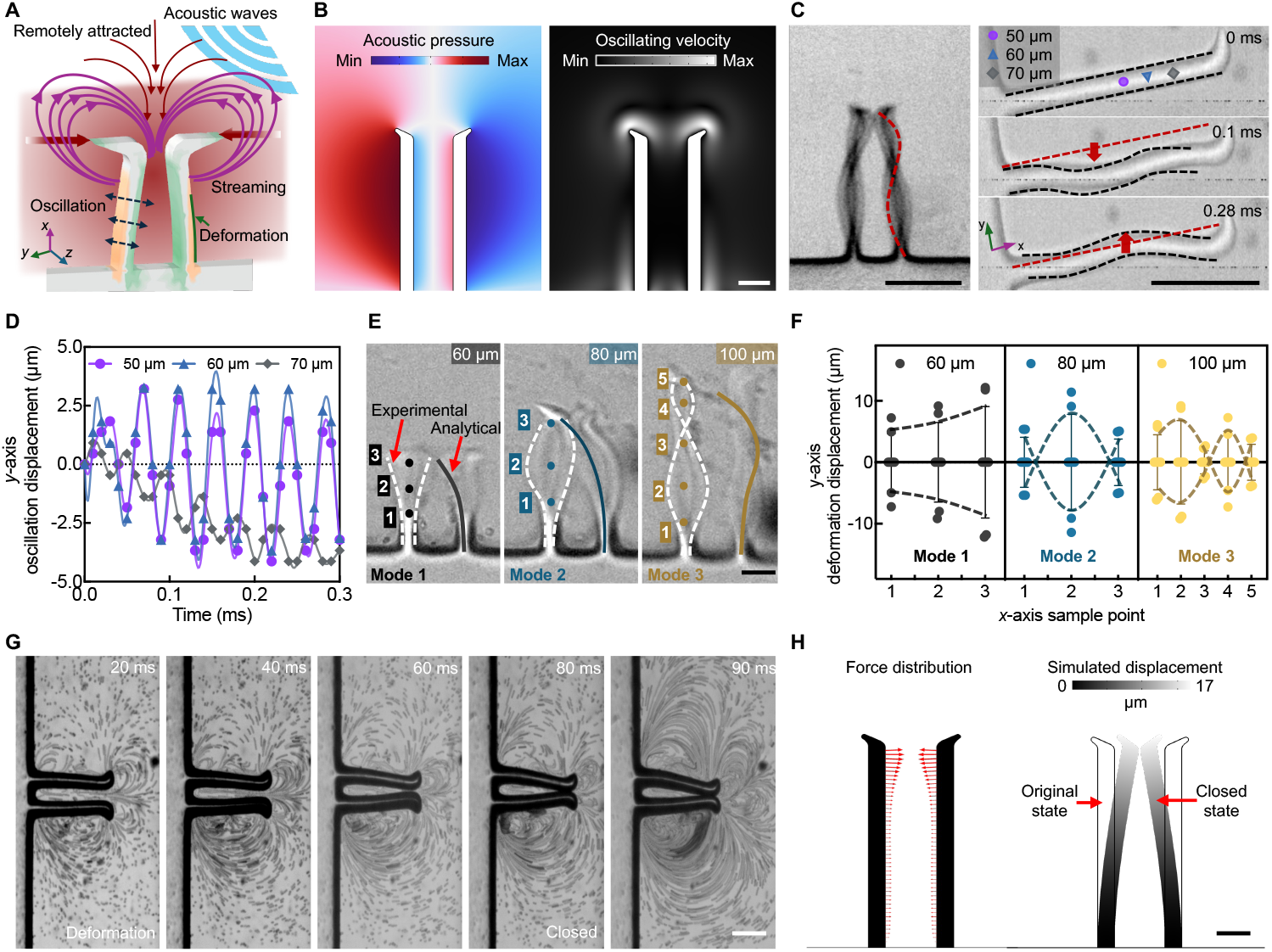
Interaction between acoustic excitations and SonoGrippers. (**A**) Schematic of the SonoGripper arm’s oscillation, deformation, induced microstreaming, and attracting force under ultrasound exposure. (**B**) Simulation of acoustic pressure (left) and oscillating velocity (right) profiles around the SonoGripper (100 µm in length, 10 µm in width, 20 µm in thickness). (**C**) Optical microscope image and time-lapse images of SonoGripper’s single arm oscillation corresponding to (B) under an acoustic excitation with a voltage of 1.2 V_PP_ and a frequency of 5.6 kHz. (**D**) Tracked oscillation displacement along *y*-axis over time at three different sample positions along *x*-axis: 50, 60, and 70 μm away from the bulky body, denoted as circle, triangle, rhombus in (C). (**E**) Different deformation modes exhibited by SonoGrippers with different arm lengths: 60, 80, and 100 μm. (**F**) Tracked deformation displacement along *y*-axis corresponding to the *x*-axis sample positions indicated by the indexes in (E), showing the different deformation modes exhibited by SonoGrippers with different arm lengths. (**G**) Time-lapse image sequence illustrates the generation of acoustic microstreaming vortices around the SonoGripper (120 µm in length, 13 µm in width, 20 µm in thickness) and stable closure. The gripper was under an acoustic excitation with a voltage of 10 V_PP_ and a frequency of 93.5 kHz. (**H**) Numerical simulation showing the forces distributed on the SonoGripper’s arms (left) and arms’ displacement profile (right) corresponding to the SonoGripper triggered by ultrasound for 90 ms in (G). Scale bar is 20 µm in (B, E, and H); 50 µm in (C and G).

Additionally, the SonoGrippers with different arm lengths presented different deformation modes under the same ultrasound exposure, which renders potentials of enhancing functionality in various environments and for different tasks such as object grasping, enveloping, or directional propulsion. Fig. 2E illustrates three deformation modes exhibited by SonoGrippers with arm lengths of 60, 80, 100 μm, experimentally and analytically (supplementary text, Fig. S4). These three SonoGrippers were attached to the same bulk body and spaced sufficiently apart to eliminate interaction effects. For characterization clarity, we defined the deformation modes based on the numbers of local undulations observed on the SonoGripper arm, which is denoted as *n*. Thus, SonoGrippers in Fig. 2E exhibited deformation mode 1, mode 2, and mode 3, respectively. Quantitative deformation displacements corresponding to the positions indicated by the indexes in Fig. 2E, were presented in Fig. 2F. In Fig. 2E, circle dots indicate the sampled positions, while dashed curves depict the deformation modes for the three SonoGrippers achieved from experiments and solid curves present that achieved from numerical analysis. These results highlight that different arm lengths produce distinct deformation modes and importantly, longer arms with higher deformation mode exhibited slightly lower maximum deformation displacement.

The microstreaming induced by acoustic wave scattering at outward-pointed sharp edges ensure a stable closure, shown experimentally in Fig. 2G and numerically in Fig. 2H. The SonoGripper was actuated using an ultrasound excitation with a driving voltage of 10 V_PP_ and a frequency of 93.5 kHz. As the arm closed, the SonoGripper reached an equilibrium and remained stable.

### Ultrafast and controllable closure of SonoGripper

When immersed in an aqueous environment and subjected to an acoustic excitation, the arm oscillation, deformation, and acoustic microstreaming around sharp edges of SonoGripper induce its millisecond-scale closure (Fig. 3A). The closure speeds can be tuned by adjusting geometrical dimensions of SonoGripper such as arm lengths and the acoustic excitation parameters such as voltages applied to the piezoelectric transducer. In Fig. 3A, the arm length and the tip-to-tip distance of SonoGripper are denoted as *L* and *D*_*TT*_, respectively.

**Fig. 3.**
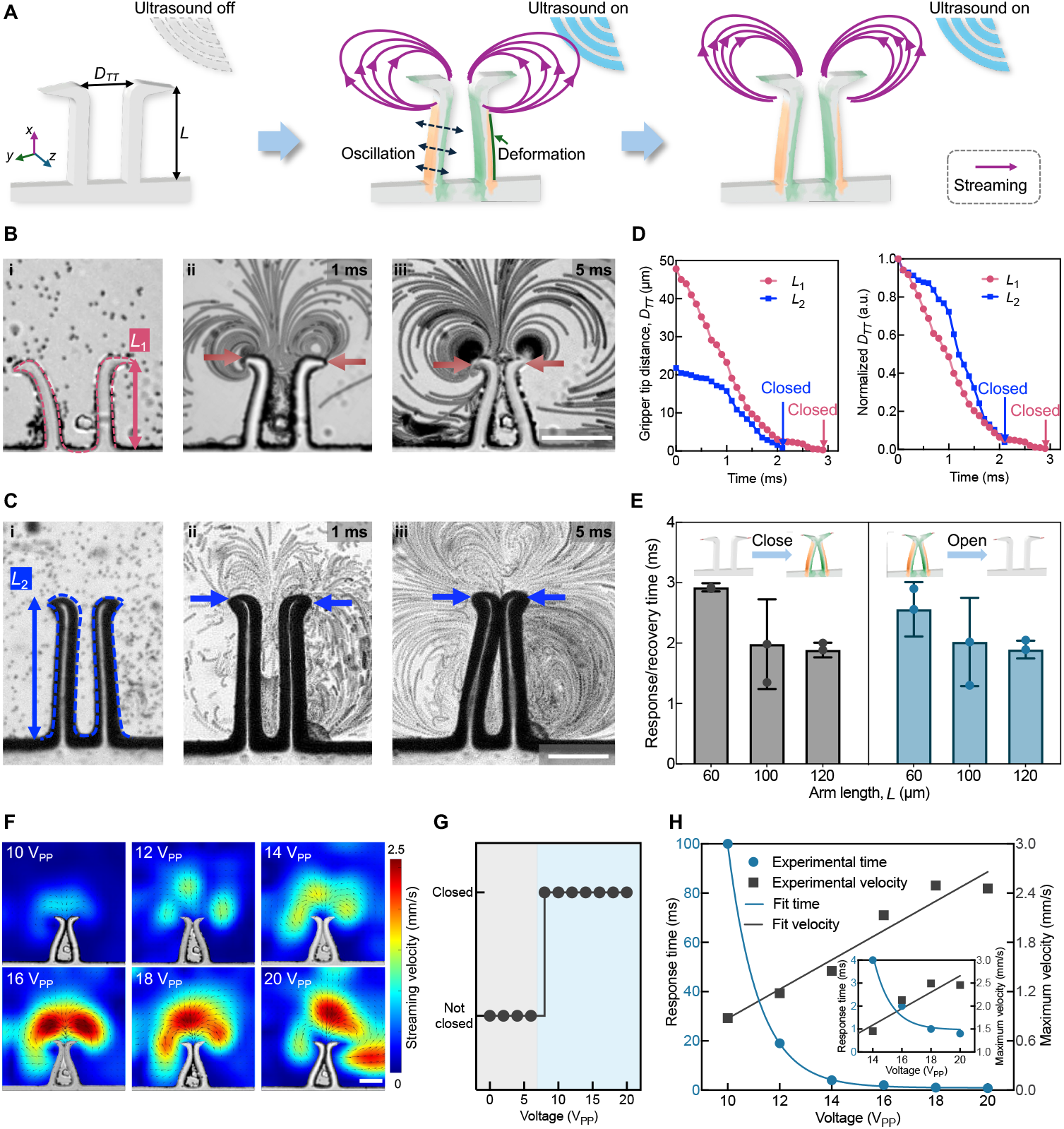
Characterization of SonoGripper closure. (**A**) Schematic illustration of SonoGripper’s closure process under ultrasound exposure. *L*: arm length, *D*_*TT*_: arm’s tip-to-tip distance. (**B, C**) Optical microscope images and high-speed camera image superpositions processed using ImageJ showing the streaming and closure behavior of SonoGrippers with arm lengths of 60 μm and 120 μm, respectively. They were triggered by ultrasound at 92 kHz and 93.5 kHz with a fixed 20 V_PP_. (i) initial state, (ii) pre-closure state, and (iii) post-closure state. (**D**) Temporal evolution of SonoGripper arm’s tip-to-tip distance *D*_*TT*_ and normalized *D*_*TT*_ (relative to the initial arm distance) illustrating closure within 3 milliseconds for the 60 μm arm length and 2 milliseconds for the 120 μm arm length. (**E**) Response and recovery time of the SonoGrippers under ultrasound stimuli, measured through fast repeated closing and opening experiments on three different SonoGrippers with arm lengths of 60, 100 and 120 μm. Results demonstrate that longer arm lengths correlate with faster response times, with a minimum response time of less than 2 milliseconds. (**F**) PIV analysis of the SonoGripper generated microstreaming under ultrasound voltages from 10 to 20 V_PP_ with a fixed frequency of 92 kHz. Arrow size represents streaming velocity magnitude, while arrow direction indicates streaming flow direction. (**G**) Phase map of SonoGripper closure as a function of ultrasound voltage, identifying a threshold voltage of 7 V_PP_ required to initiate closure. (**H**) Plots correlating ultrasound voltage with SonoGripper closing time and maximum velocity along the *x*-axis above SonoGrippers. Higher ultrasound voltages result in faster closing times and higher maximum streaming velocities (SonoGripper dimensions: 20 μm thickness, 90 μm arm length, and 6 μm initial arm tip-to-tip distance). Scale bar is 50 µm.

To characterize the geometric effects on the closure behavior, we fabricated SonoGrippers with varying *L*, via specific photomask designs and UV light exposure times. Then, we triggered these SonoGrippers under acoustic excitation generated by the PZT experimental setup. Fig. 3B and 3C present two SonoGrippers with arm lengths of 60 µm and 120 µm, respectively actuated at frequencies 92 kHz and 93.5 kHz with the same voltage of 20 V_PP_. The dispersed tracer particles’ superposed trajectories (see Fig. 3B ii, iii, Fig. 3C ii, iii, and Fig. S5, Movie S2) shows that after closure, the acoustic microstreaming vortices contracted and moved, while an additional vortex emerges outside the arms (Fig. 3C iii). Simultaneously, streaming velocity increased above the gripper’s center, while flow velocity within the closed arms dropped to near zero (see Fig. S5). These results highlight a complex nonlinear interaction between SonoGripper and acoustic microstreaming, significantly influenced by arm lengths and arm tips’ relative position. We further tracked the *D*_*TT*_ over time, and normalized *D*_*TT*_ against the initial arm separation distance for these two SonoGrippers with varying arm lengths, Fig 3D. The closure times of SonoGrippers with 60 µm and 120 µm arm lengths were measured to be 3 ms and 2 ms, respectively. Additional closure trials depict the stability of SonoGripper triggered by ultrasound (Movie S3). Upon deactivating the transducer, the acoustic streaming ceased instantaneously due to the gripper’s deformation in the elastic regime, allowing the gripping arms to fully return to their original state within 3 ms, as shown on the right panel of Fig. 3E, in Fig. S6A and Movie S4. Fast repeated closing and opening experiments through switching on and off acoustic actuation demonstrated that SonoGrippers with longer arms exhibited faster response times (Fig. 3E). A representative case of a SonoGripper with 100 µm arm length undergoing repeated actuation in response to the acoustic signal (Fig. S6B and Movie S4) confirms the fast response times and consistency of repeated operation. These ultrafast closing and opening times, which can be tunable by arm lengths, enabled high-speed, repeatable, controllable gripping cycles, allowing for multiple grips per second simply by toggling the acoustic actuation or applying pulsed ultrasound stimulation. Meanwhile, the repeated closing and opening experiments also validated the durability of SonoGrippers. Additionally, the initial arm separation distance was found to influence closure dynamics, further supporting the tunability of the SonoGripper’s closure through geometric design adjustments to accommodate diverse gripping requirements (Fig. S7).

To further investigate the voltage-influenced closure behavior, we examined acoustic microstreaming profiles, that affect suction forces inside SonoGripper and net inward force acting on the SonoGripper arms, at varying ultrasound voltages. High-speed imaging and PIV analysis quantified streaming velocity distributions around the SonoGripper. Fig. 3F presents PIV-derived streaming velocity under voltages ranging from 10 to 20 V_PP_, showing that higher voltages enhance streaming velocities, particularly around the SonoGripper arms, thereby showing the potential to expand the remote attraction and gripping region. Additionally, Fig. 3G identifies a voltage threshold of 7 V_PP_ required to initiate closure. Below this threshold, no observable shape transformation of SonoGripper arms since the acoustic energy was insufficient to overcome the elastic strain resistance of the hydrogel. Beyond this threshold, increasing the voltage further accelerated closure, exhibiting an approximately exponential decay of closure time and voltage (Fig. 3H). Additional closure time measurements for three separate trials with different acoustic voltages further validated these results (Fig. S8). Simultaneously, the maximum streaming velocity along *x*-axis above the SonoGrippers increased linearly with the applied voltage (Fig. 3H). These results confirm that stronger acoustic actuation reduces closure time and enhances acoustic streaming velocity, thereby improving remote object attraction and enabling precise control over closure speed and gripping strength through voltage modulation.

### Remote, and multidirectional attraction and gripping of diverse objects

The acoustic-soft matter interactions endow SonoGripper with dexterous attraction and gripping capabilities of diverse objects. First, we characterized its ability to attract and grip objects from multiple directions and measured its effective attractable regions, as schematically illustrated in the center of Fig. 4A. The attracting and gripping processes were analyzed by tracking object trajectories until they reached the interior of the SonoGripper using high-speed imaging and TrackMate in ImageJ. The magenta lines and magenta circles in Fig. 4A show the multidirectionally attracted object trajectories. Objects approaching from the left, right, near above, and far above directions were successfully attracted, demonstrating the broad attractable range and multidirectional manipulation capability of the SonoGripper. To further quantify the attracting and gripping processes, we measured the distance evolution over time for diverse objects originating from left, right, and above directions, respectively (Fig. 4B). The residual trajectory distance (*d*_*t*_) defined as the remaining distance to the gripped site at interior of the SonoGripper was tracked across multiple trials. Results show that initial distances ranged from 50 to 500 µm (0.625-6.25 times the SonoGripper’s body length), with an effective attraction distance as the crow flies exceeding 400 µm (5.0 times the SonoGripper’s body length). Across all cases, gripping from left and right was completed within 150 ms, while gripping from above was completed within 50 ms, highlighting the ultrafast, remote and multidirectional attraction capability of SonoGripper. The fast attraction trajectories from multiple directions are consistent with the acoustic streaming vortices around SonoGripper, further validating the role of acoustic microstreaming in remote object attracting and gripping. Then, we accessed the remote attractable region sizes in relation to different target volumes, Fig. 4C. As the target volume increased, the radius of the attractable region (*r*) was observed to increase accordingly. This positive correlation confirms that larger targets with increased surface area-to-volume ratios interact more effectively with acoustic streaming vortices, leading to stronger attraction forces and an expanded attractable region. The results also reveal the capability of SonoGripper to attract and grip multiscale objects.

**Fig. 4.**
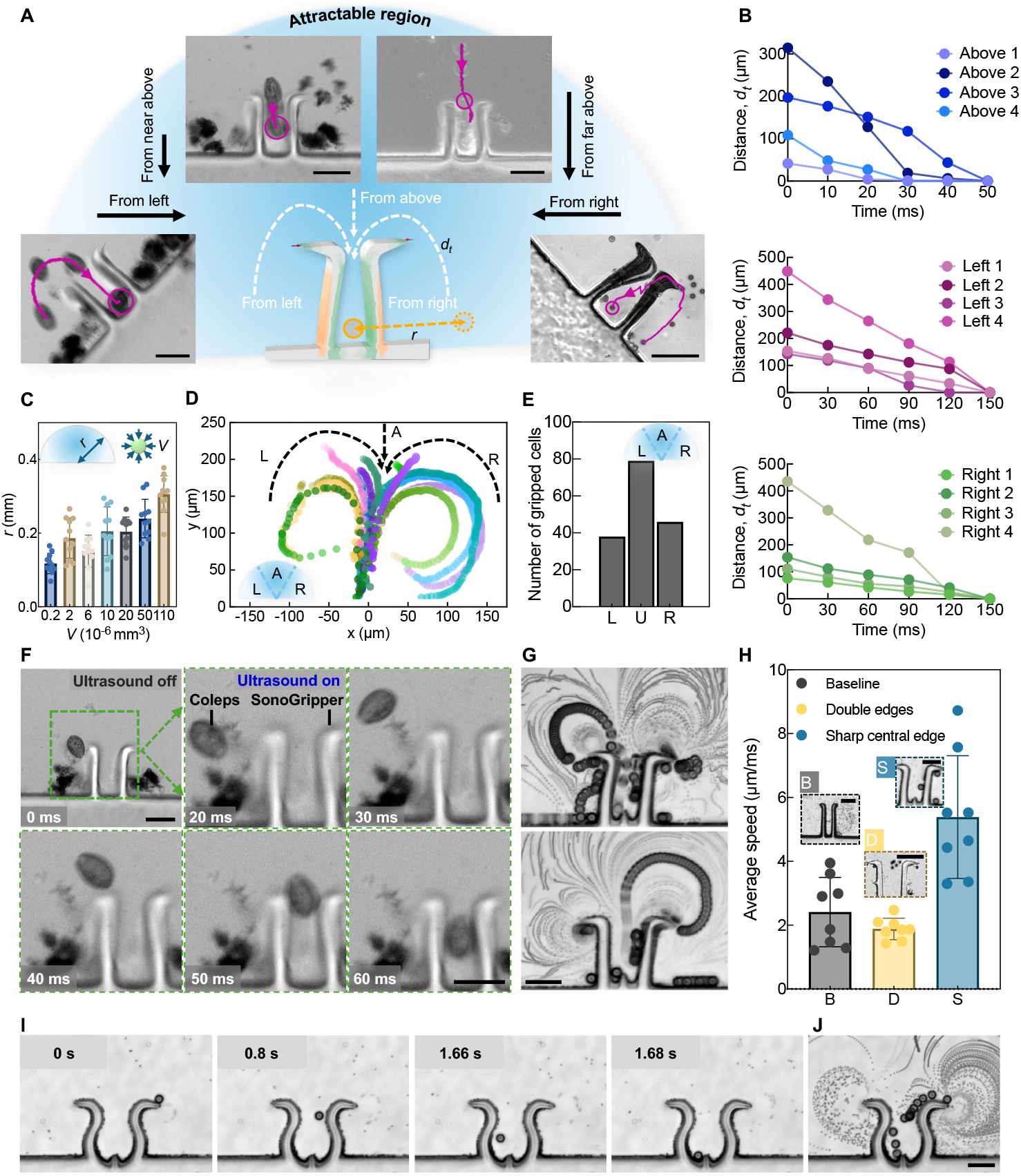
Attraction and gripping of objects using SonoGripper. (**A**) Schematic illustration of SonoGripper attracting and gripping objects from different directions (left, above, and right). The blue background highlights the attractable region, where *r* denotes the radius of attractable region, and *d*_*t*_ represents the residual distance as objects approach the interior of SonoGripper from various directions. The microscope image superpositions with object trajectories analyzed via ImageJ present that SonoGripper remotely and multidirectionally attract and grip individual targets. (**B**) Evolution of *d*_*t*_ over time for various objects approaching from the above, left, and right directions, respectively. (**C**) Characterization of attractable region size relative to different object volumes. (**D**) Tracked trajectories of yeast cells simultaneously attracted from different directions, with transparency gradients indicating time evolution. The directions are labeled by: L (Left), A (Above), and R (Right). (**E**) Number of yeast cells gripped from different directions and categorized by directions. (**F**) Fluorescent images showing a 20 µm Coleps gripped by the SonoGripper within 60 milliseconds. (**G**) Microscope image superposition showing the SonoGripper with a sharp central edge attracting and gripping 15 μm particles from both left and right sides in a 2 μm bead solution under a 94 kHz and 17 Vpp acoustic excitation. (**H**) Comparison of average gripping speeds among the baseline SonoGripper and SonoGripper variants featuring double edges at the arm tips or a sharp central edge. (**I**) Image sequence depicting a SonoGripper with curved arms and a sharp central edge attracting and gripping 15 μm particles in a 2 μm bead solution under a 94 kHz and 7 V_PP_ acoustic excitation. (**J**) Superposition image corresponding to (I) illustrating the trajectory of an attracted 15 μm particle. Scale bar is 50 μm.

We next evaluated the SonoGripper’s ability to grip multiple objects continuously by dispersing yeast cells around the gripper and recording their movement under acoustic actuation. Using the TrackMate plugin in ImageJ, we visualized yeast cell trajectories before SonoGripper closure, confirming continuous multidirectional gripping. Fig. 4D presents multiple yeast cell paths attracted from different directions, with transparency gradients indicating temporal evolution. These trajectories perfectly align with the acoustic streaming vortices, underscoring the critical role of microstreaming profiles in remote object attraction and defining the attractable region. To further quantify continuous multidirectional gripping, we categorized the attracted yeast cells based on their approach direction (left, above, and right), Fig. 4E. Results revealed that gripping from the above direction was dominant, consistent with PIV-derived velocity maps, which showed maximum streaming velocity above SonoGripper’s center. This phenomenon arises due to flow superposition from the left and right sharp edges, reinforcing the role of acoustic streaming dynamics in shaping the effective gripping region. These findings highlight SonoGripper’s potential for high-throughput manipulation, enabling simultaneous capture of multiple microscale objects.

To assess the SonoGripper’s adaptability, we tested its performance on biological samples and living microswimmers of varying sizes, shapes, and motilities. We examined Coleps, a highly motile microswimmer, by suspending them in a drop of deionized water and actuating the SonoGripper using ultrasound. Fluorescence imaging sequences captured the gripping process of a 20-µm Coleps within 60 ms (Fig. 4F), demonstrating the SonoGripper’s ability to manipulate dynamic targets. The incomplete grip observed in this case was attributed to the size of Coleps, which can be addressed by designing longer arms. Additional tests confirmed successful gripping of yeast cells and Human Embryonic Kidney (HEK) cells with sizes of 3-20 μm (Fig. S9 and Movie S5), underscoring the SonoGripper’s broad adaptability across diverse biological systems.

Finally, we explored the effects of additional microstructures on acoustic-soft matter interaction and gripping behavior of SonoGrippers, whereby optimizing gripping efficiency. We fabricated SonoGripper variants with distinct microstructural modifications, including double edges at arm tips, a sharp central edge, and curved arms with a sharp central edge. Microscope image superimpositions in Fig. 4G illustrate the performance of a sharp central edge variant, which successfully attracted and gripped 15 µm particles from both the left and right directions under a 94 kHz, 17 V_PP_ acoustic excitation. To quantitatively assess the impact of these structural modifications, we analyzed the attraction and gripping speeds across different SonoGripper designs (Fig. 4H, Fig. S10, Fig. S11, Movie S6). Compared to the baseline SonoGripper without any microstructures, the speed improvements were yielded by the sharp central edge variant, which were attributed from that the sharp central edge significantly enhanced localized acoustic streaming, in turn, creating stronger focused forces and resulting in faster gripping speeds. In contrast, double edges at the arm tips slightly reduced gripping speed, despite creating additional vortices in the acoustic streaming profile. This reduction might be attributed to the complex flow patterns generated by the double edges, which could disrupt the coherence of the acoustic streaming and lead to less forces. Additionally, Fig. 4I presents the effective attracting performance of a curved arm SonoGripper with a sharp central edge. The temporal image sequence demonstrated the attraction and gripping of 15 µm beads, with superimposed trajectories (Fig. 4J) depicting the captured bead’s pathway. These findings confirm the significant role of acoustic streaming in SonoGripper performance and highlight the importance of microstructural engineering in refining the gripping mechanism for enhanced precision and efficiency.

### Mechanism of SonoGripper

As schematically illustrated in Fig. 5, we take one arm of the SonoGripper as a slender, soft hydrogel beam (with length *L*, height *h*, and width *b*), clamped at one end and free at the other, undergoing transverse deflection *w*(*x, t*) in response to acoustic excitation and fluid interaction. Under moderate strain, it exhibits strongly nonlinear stress-strain behavior, which motivates using a nonlinear beam theory. At ∼4 kHz, the acoustic wavelength in water (∼0.37 m) is typically much larger than the beam, so we can treat the incident pressure as nearly uniform over the beam. The beam is submerged in water, and a sharp edge at the free tip induces acoustic streaming, which can be modeled as static force free of time. We model the beam as slender and adopt Euler–Bernoulli kinematics (*63*) (plane sections remain normal), the governing equation, accounting for elastic bending, effective inertia (including added mass), viscous damping, and streaming-induced forcing, is derived in the Supplementary Material and, for small deflection, approaches:

**Fig. 5.**
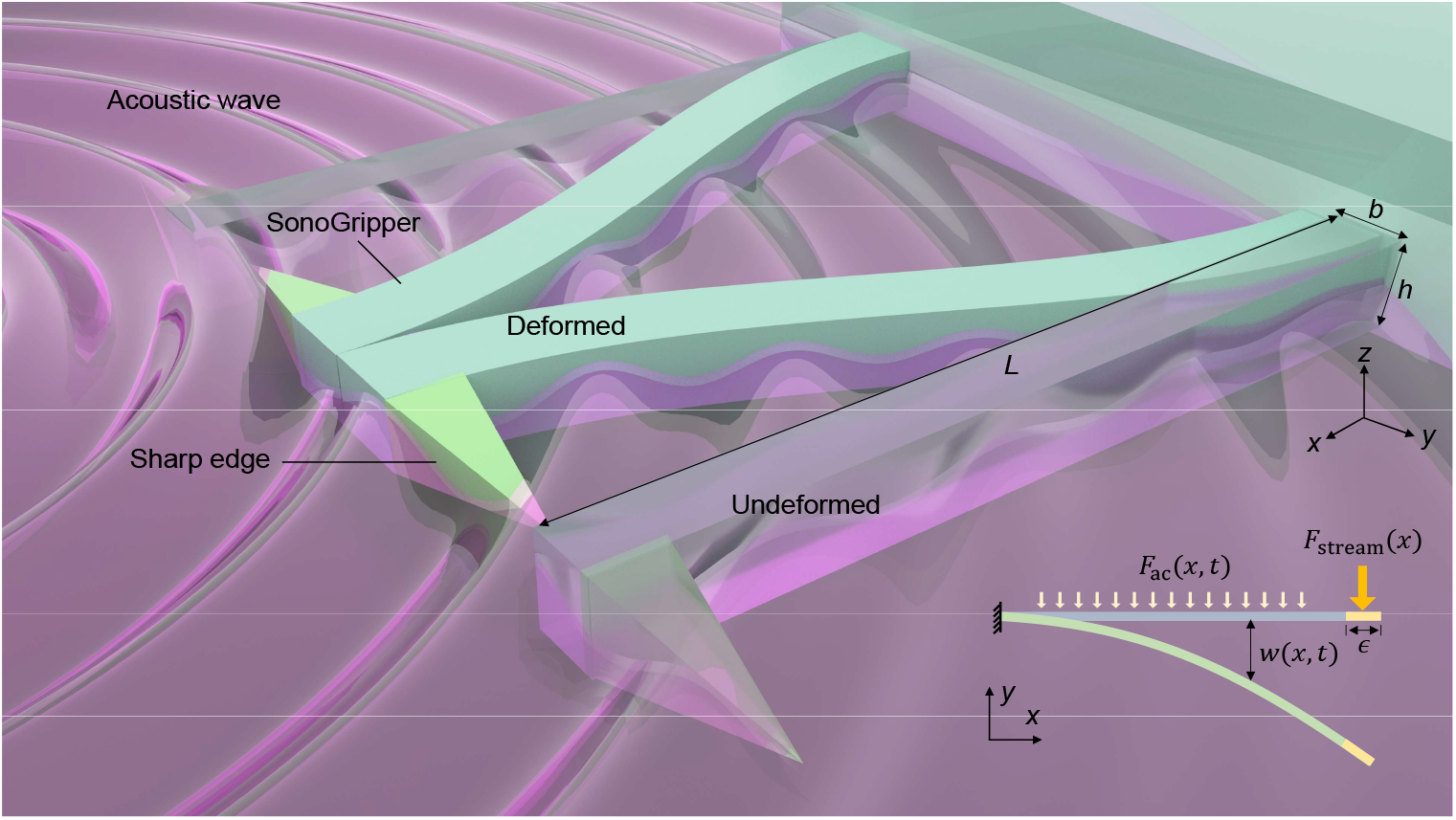
Modeling of the SonoGripper’s deformation under the ultrasound excitation. The SonoGripper, with length *L*, height *h*, and width *b*, undergoes deformation under acoustic excitation. The zoomed-in schematic (right bottom corner) illustrates a simplified two-dimensional model, where the SonoGripper body is subjected to a time varying uniformly distributed acoustic force *F*_ac_ (*x, t*). The concentrated acoustic force acting near the sharp edge is approximated as a constant point load *F*_stream_(*x*) with an effective small loading width *ϵ*. The time-dependent transverse deformation of the beam at position *x* is denoted by *w*(*x, t*).

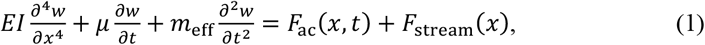

where *E* and *I* are the Young’s modulus and second moment of area of the beam, *μ* is the hydrodynamic damping per unit length, *m*_eff_ is the effective mass per unit length, including the added mass due to the fluid-structure interaction, *F*_*ac*_(*x, t*) is the harmonic acoustic pressure force per unit length, and *F*_stream_(*x*) is the steady force due to the acoustic streaming force working pointwise (*ϵ* ≪ *L*) at the tip of the free end.

We employed a mode superposition approach to model the deformation of cantilevered soft hydrogel beams under acoustic excitation. The beam equation was discretized using finite differences, and an eigenvalue analysis was performed to extract mode shapes and natural frequencies. Harmonic excitation was modeled as a sinusoidal tip force, while acoustic streaming was treated as a static load at the free end; both were projected onto the modal basis to determine the modal amplitudes. The full spatiotemporal deformation was reconstructed by summing the weighted mode shapes. This linear framework assumes small deformations, linear elasticity, and linearized hydrodynamic loading, an appropriate approximation for soft hydrogels in moderate underwater acoustic fields. The resulting frequency response function captures both dynamic and static deflection components:

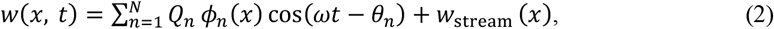

where *ϕ*_*n*_ is the *n*-th mode shape and *Q*_*n*_ corresponding amplitude. *ω* denotes the angular frequency of excitation, and *θ*_*n*_ is the phase shift associated with the *n*-th mode. The contribution *w*_stream_(*x*) represents the deformation induced by the acoustic streaming force (see details in Supplementary Materials). The model is intended to capture the essential features of the closure and oscillation behavior of the SonoGripper, including the spatially distributed, time-varying deformation along the beam. Importantly, while the first bending mode dominates under low-frequency excitation, higher-order modes become notable for longer beams or under higher-frequency acoustic actuation. These higher modes introduce localized curvature, enabling complex shape transformations such as S-bends, multi-peak undulations, or tip curling, functions critical for tasks like object grasping, enveloping, or directional propulsion. By tailoring the acoustic input and beam geometry, the modal content can be selectively activated, effectively programming the SonoGripper’s deformation profile for multifunctional performance.

## Discussion

Drawing inspiration from Venus flytrap and *Vorticella*, this study presents a microscale soft SonoGripper fabricated from polyethylene glycol diacrylate (PEGDA) hydrogel, capable of acoustically driven oscillation and deformation as well as of attracting and gripping objects. Compared with conventional microgrippers, our SonoGripper offers soft, deformable structures that enable flexible shape transformations while minimizing the risk of sample damage. Its microscale design allows for operation in hard-to-reach environments, making it well-suited for delicate and minimally invasive applications. Additionally, the custom-built one-step photopolymerization technique used to fabricate SonoGripper facilitates the scalable fabrication with varied geometries and additional microstructural features, underscoring the simplicity and versatility of the manufacturing process.

The mechanism of SonoGripper is governed by the interaction between acoustic actuation and soft structures, which is not yet fully understood especially at the microscale. Upon acoustic exposure, the soft SonoGripper arms induce rapid oscillation and deformation, generating strong localized streaming flows around sharp edges at the arm tips that attract samples and initiate closure. The deformation modes, velocity and profile of the streaming flow play a significant role in the attracting and gripping behaviors. This mechanism enables fast gripping, overcoming the limitations of swelling-based hydrogel actuators, which often require minutes to achieve deformation. High-speed imaging confirmed a closing time of only 2 milliseconds, and the SonoGripper immediately returned to its original state when the acoustic actuation ceased, enabling repeated gripping operations within a second through toggled or pulsed ultrasound actuation. Tunable deformation modes also enhance SonoGripper’s functionality in viscous environments. Our simulations and theoretical studies further confirmed this fundamental mechanism of SonoGripper. Moreover, unlike mechanisms that rely on temperature or pH changes, this work leverages a biocompatible, untethered structure with harmless acoustic stimulation, making it highly suitable for in vivo applications.

Systematic characterization demonstrated that geometrical dimension tuning and acoustic excitation modulation enable precise and controllable gripping performance. Increasing the arm length or actuation voltage reduced the response times, while higher voltages also amplified streaming velocities, particularly above the SonoGripper. Moreover, the SonoGripper successfully manipulated objects from multiple directions (above, left, and right), both individually and simultaneously, highlighting its adaptability for complex and high-throughput micromanipulation tasks. Sample trajectory tracking confirmed that acoustic streaming profiles influence attracting and gripping behavior, suggesting that real-time tuning of streaming patterns could further enhance gripping precision and flexibility. The demonstrated capability to grip microparticles, living cells, and motile microorganisms positions the SonoGripper as a promising platform for applications in targeted drug delivery, cell manipulation, microassembly, and environmental monitoring. Additionally, incorporating microstructural modifications, such as double edges at the arm tips, sharp central edges, and curved arms, allowed for localized control over streaming-induced forces, further altering and refining the gripping performance.

Despite these advances, the study highlights complex interactions between soft matter and acoustic excitation, which introduce complexity in predicting and controlling gripping behavior. The relationship between SonoGripper’s properties (like geometrical parameters and stiffness), interaction to acoustic excitations, and gripping efficiency requires further investigation to develop predictive models that can guide design and dynamic acoustic optimization for specific applications. Additionally, the integration of microstructural features introduced complex flow patterns, which could either enhance or disrupt localized acoustic streaming and therefore influence the gripping efficiency. Future research should focus on systematic microstructural optimization to maximize the gripping efficiency.

These findings establish the SonoGripper as a powerful microscale gripping technology, integrating fast controllable actuation, biocompatibility, untethered operation, and versatile object handling. It paves the way for advancements across multiple scientific domains. From a physics and acoustofluidics perspective, the SonoGripper provides an exciting platform to explore acoustic interactions with soft matter at the microscale. In the field of microdevices and microfluidics, integrating the SonoGripper into lab-on-a-chip platforms could enable precise, high-throughput sample manipulation for biological assays, drug screening, and single-cell studies. Furthermore, in flexible electronics, SonoGripper could play a key role in handling and assembling soft electronic components with minimal mechanical strain. In biomedicine, SonoGripper’s acoustic excitation mechanism is particularly promising, since acoustic waves penetrate biological tissues with minimal invasiveness. Future work should focus on further refining microstructural designs, dynamically tuning acoustic excitations, exploring the potential to dynamically adapt and navigate through complex vascular networks and expanding applications in biomedical and microengineering domains. Addressing them would pave the way for next-generation microscale gripper technologies.

## Materials and Methods

### Experimental Design

The primary objective of this study was to explore the acoustic-soft matter interaction and develop a soft microscale SonoGripper capable of ultrafast and programmable response to the acoustic actuation. SonoGrippers were fabricated using a one-step photopolymerization technique with polyethylene glycol diacrylate (PEGDA) hydrogel. Geometrical properties, such as arm length and width, were systematically varied, and microstructural features, such as double edges and sharp central edges, were incorporated to optimize performance. A piezoelectric transducer (PZT) operating at frequencies ranging from 5 to 100 kHz with a voltage of 1-20 V_PP_ actuated the SonoGrippers. High-speed microscopy, particle image velocimetry (PIV), and trajectory tracked by ImageJ were employed to characterize deformation, oscillation, acoustic streaming, and gripping behavior under the acoustic excitation. The response time and controllability by gripper’s structure designs and actuation parameter tuning were characterized. Also, the SonoGripper’s ability to attract and grip objects was tested using diverse samples including yeast cells, HEK cells and motile microorganisms (e.g., Coleps). SonoGripper variants with additional microstructures were fabricated and tested to assess their impact on interaction with ultrasound and SonoGripper’s behavior.

To ensure reproducibility and minimize bias, the prespecified components include photosensitive hydrogel mixture, photomask preparation, photopolymerization process, experimental system setup, sample preparation, statistical analysis methods and numerical simulations.

### Photosensitive Hydrogel Mixture

The photosensitive hydrogel mixture used to fabricate the soft SonoGripper consisted of a 5 mL synthetic solution containing 85 vol% polyethylene glycol diacrylate (PEGDA, molecular weight of 700, Sigma-Aldrich) and 15 vol% photo-initiator (2-hydroxy-2-methyl1-phenyl-propan-1-one, Darocur 1173, Sigma-Aldrich). The photo-initiator triggers the crosslinking of PEGDA 700 upon exposure to ultraviolet (UV) light (Nikon Intensilight C-HFGI, intensity: 130 W). To facilitate alignment of the photomask and visualization of the SonoGrippers during the polymerization fabrication process, three droplets (∼90 μL) of blue dye (Trawosa) or Rhodamine B solution (Sigma-Aldrich) were added to the 5 mL solution, rendering the mixture fluorescent.

### Photomask Preparation

The photomask used for fabricating the SonoGrippers were designed using AutoCAD (Autodesk, Inc., San Rafael, CA) software. The AutoCAD designs were printed at a high-resolution of 20,000 dpi by CAD/Art Services, California, ensuring accurate reproduction of fine details during photopolymerization. The bitmap images with transparent and opaque regions were printed onto films using a high precision photoplotter, creating photomasks with well-defined patterns. The prepared photomasks enabled the rapid and scalable production of SonoGrippers with precise control over their dimensions and structural features, laying the foundation for systematic characterization and optimization of their gripping performance.

### Photopolymerization based Fabrication Method

Our custom-built UV-light photopolymerization setup is developed on an inverted microscope (NIKON, Eclipse Ti). A UV lamp (Nikon Intensilight C-HFGI) with an intensity of 130 W and a shutter controller (Vincent Associates, VCM-D1) controlled via LabView are connected to irradiate a high-resolution photomask (CAD/Art Services, Inc.) inserted into the field stop of the microscope. A ∼10 μL droplet of the photosensitive hydrogel mixture was dispensed onto the fabrication area of a glass substrate (60 mm × 20 mm × 0.13 mm). The dispensed photosensitive hydrogel mixture was then spread by placing a round coverslip (22 mm diameter, 0.15 mm thickness) to achieve a uniform thickness of approximately 20-25 µm. When the photomask was irradiated, the UV light passed through photomask and was focused by a 20 × objective lens, polymerizing the photosensitive hydrogel mixture. The photomask patterned the UV light, and the 20 × objective lens reduced the structure size by a factor of 16.5 ×. Depending on the desired geometry of SonoGrippers, the exposure time varied between 20 ms and 120 ms. Notably, insufficient UV exposure time may result in incomplete polymerization or excessively soft structures that break under high excitation voltages, while excessive exposure may lead to overly thick structures that fail to oscillate effectively under acoustic excitations. After fabrication, the coverslip was lifted. Finally, the SonoGrippers were cleaned with isopropanol (IPA) and deionized water to remove residual mixtures and contaminants. Thereby, one SonoGripper was produced per photomask pattern. By moving the microscope stage with the mounted glass substrate over the objective lens, multiple SonoGrippers could be fabricated rapidly.

### Experimental System Setup

The experimental setup was constructed on a 25 mm × 75 mm × 1 mm microscope glass substrate (Menzel-Gläser). A circular piezoelectric transducer (27 mm diameter, 0.54 mm thickness, resonance frequency 4.6 kHz ± 4%, Murata 7BB-27-4L0) was affixed to the glass substrate using a two-component epoxy adhesive (2-K-Epoxidkleber, UHU Schnellfest). A droplet deionized water (50-80 µL) was dispensed onto the fabricated SonoGripper laying on the substrate and covered with a coverslip to create a thin and stable aqueous environment with a thickness of 90 - 150 µm. The piezoelectric transducer was connected to an electronic function generator (AFG-3011C, Tektronix), which was further connected to a signal amplifier (0-60 V_PP_, 15 × amplification, High Wave 3.2, HF-Amplifier, Digitum-Elektronik) to generate acoustic waves with tunable excitation frequencies and voltages. The acoustic setup was then mounted on an inverted optical microscope (Eclipse Ti, Nikon or Axiovert 200M, Zeiss) for observation. Tracers (monodisperse polystyrene microspheres, 2-15 µm, Polybeads®) were used to visualize arm oscillation, deformation and acoustic microstreaming induced by soft SonoGripper triggered by ultrasound. The arm oscillation, deformation, and acoustic microstreaming were recorded using a high-speed camera (Chronos 1.4, Kron Technologies) that was integrated to the microscope.

### Sample Preparation

To assess the SonoGripper’s versatility, samples with multiple sizes, shapes, and motilities were tested, including polystyrene microparticles (15 µm diameter, microsphere), yeast cells (3∼7 µm, typically sphere or ellipsoid), Human Embryonic Kidney (HEK) cells (3∼20 µm, round or elongated), and motile microorganisms such as Coleps (20 µm diameter, Barrel-shaped). Yeast cells were cultured and dispersed in deionized water to achieve a uniform solution, while Coleps were collected from pond water and concentrated to ensure sufficient density for testing. HEK cells were cultivated, kindly provided by the Institute of Pharmacology and Toxicology for the University of Zurich, and suspended in cellular media (DMEM + GlutaMAX (Gibco), supplemented with 5% (v/v) FBS). A ∼30 µL droplet of sample solution was placed on a glass slide with the SonoGripper, and their behavior under acoustic actuation was recorded using high-speed camera and fluorescence microscopy.

### Statistical Analysis

The SonoGripper behavior under ultrasound exposure was recorded using a high-speed camera (Chronos 1.4, Kron Technologies) and/or a fluorescent camera (Coolsnap EZ, Photometrics) attached to the optical inverted microscope. Recording frame rates range from 50 to 32,668 frames per second (fps). The recorded footage was analyzed using ImageJ and PIVlab in MATLAB software. The Young’s modulus of hydrogel structures was measured using atomic force microscope (AFM, Dimension FastScan®).

### Numerical Simulations

Numerical simulations were conducted using COMSOL Multiphysics finite element software by adapting our recently reported numerical model (*64*) the SonoGripper geometry. The acoustic streaming effect was modeled using a perturbation expansion method, where flow variables were decomposed into first- and second-order components. The first-order terms, representing the system’s acoustic response, were treated as harmonic oscillations matching the actuation frequency. In contrast, the second-order terms captured the system’s time-averaged behavior, varying over a longer timescale than the acoustic wave period. This decomposition allowed the Navier-Stokes equations to be split into two distinct, linear sets of equations—one for each order of the flow variables. These were solved sequentially: the first-order equations were addressed in the frequency domain, while the second-order equations were solved as steady-state problems.

2D simulations were conducted on an overall rectangular geometry comprising the fluid domain and the gripper domain. Following our prior work (*64*), a background field was prescribed to model the acoustic actuation. The fluid-solid interface was prescribed velocity and traction continuity boundary conditions. The domain was discretized with an unstructured triangular mesh. For the first-order problem, pressure and velocity are approximated using P2-P3 composite elements, where P2 and P3 correspond to second- and third-order Lagrange polynomial triangular elements, respectively. Similarly, for the second-order problem, pressure and velocity are approximated using P1-P2 elements. A direct solver was used to compute solutions for both sets of equations.

## Acknowledgments

This project has received funding from the European Research Council under the European Union’s Horizon 2020 Research and Innovation Programme (grant agreement no. 853309, SONOBOTS); the Swiss National Science Foundation under project funding MINT 2022 (grant agreement no. 213058) and Spark 2023 (grant agreement no. 221285); and an ETH research grant (agreement no. ETH-08 20-1). R.W. is funded by the Deutsche Forschungsgemeinschaft (DFG, German Research Foundation) — 535275785. N.N acknowledges funding support from the United States National Science Foundation (OIA-2229636, CBET-2407937).

## Author contributions

Conceptualization: D.A.

Methodology: C.Z., V.W., C.D., T.B., Z.S., D.A.

Investigation: C.Z., V.W., K.K., C.D., T.B., Z.S., J.S.

Visualization: C.Z., V.W., K.K., C.D., Z.S., J.S.

Theoretical understanding: Z.S., J.S., R.W., D.A.

Numerical simulation: K.K., Z.S., J.S., R.W., N.N.

Supervision: S.L., R.W., N.N., D.A.

Writing—original draft: C.Z., V.W., C.D., Z.S.

Writing—review & editing: Z.S., S.L., R.W., N.N., D.A.

## Data and materials availability

All data are available in the main text or the supplementary materials. Additional data generated and analyzed during this study, such as experimental results and simulation parameters that support the plots and findings, are available from the corresponding author upon reasonable request.

## Supplementary Materials

### Supplementary Text

#### S1. Young’s modulus estimation using atomic force microscope

We estimated the Young’s modulus of SonoGrippers fabricated with varying UV exposure times. The estimation was based on z-axis displacement measurements of the cantilever tip obtained using atomic force microscope (AFM) and Sneddon model (*1, 2*). AFM measurements were conducted in a Peak Force Mapping mode in Air, with a scanning size of 200 nm and a scanning rate of 1.00 Hz. The cantilever used had a spring constant of 42 N/m, a tip radius of 8 nm, and a tip opening half angle of 25 degrees. During scanning, the relationship between the applied force and the z-axial displacement of the cantilever tip was continuously monitored and captured (Fig. S2). Based on the measured force, we calculate the Young’s modulus using the Sneddon model, described by the following equation,

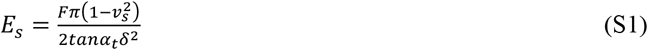

where *E*_*s*_ is the Young’s modulus of SonoGripper to be estimated, *F* is the applied force, *?*_*s*_ is the Poisson’s ratio of the SonoGripper, *α*_*t*_ is the cantilever tip’s opening half angle, and *δ* is the indentation depth of the cantilever tip. We performed measurements for SonoGrippers fabricated with different UV exposure times varying from 20 to 120 ms with an incremental step of 10 ms.

#### S2. Calculation of SonoGrippers’ Poisson’s ratio

To determine the Poisson’s ratio, we first measure the initial dimensions of the SonoGripper (*x*_0_ and *y*_0_) along the x-axis and y-axis, respectively, using recorded footage. Subsequently, we measure its dimensions after achieving stable closure (*x*_1_ and *y*_1_). The deformation is then quantified as,

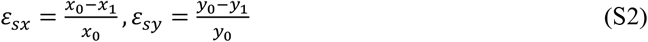

Using the measured deformation along both dimensions, the Poisson’s ratio is calculated as

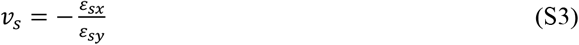

For our SonoGripper, the calculated Poisson’s ratio is 0.49.

#### S3. Deformation of the SonoGripper

We model one arm of the SonoGripper as a slender, cantilevered soft hydrogel beam of length *L* with a rectangular cross-section, clamped at one end and free at the other, undergoing linearly elastic deformation in response to an acoustic field. In the hydrodynamic model, we assume an incompressible fluid and small-amplitude velocity field. The slender beam interacts linearly with the surrounding fluid through friction and added mass. Due to its symmetric cross-section, we neglected acoustic streaming forces along its body. Owing to the small size of the beam tip, we neglect its linear hydrodynamic interactions but incorporate its nonlinear (quadratic) streaming force as a localized point force. Furthermore, we assume local fluid–structure interactions, neglecting acoustic reflections and the extension of streaming fields from the tip to other parts of the beam.

We parameterize the beam by length *x*∈[0, *L*], and describe its deformation using the angular slope *θ*(*x*). The unit tangent and normal vectors are then given by

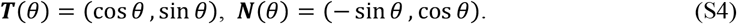

Accordingly, the position of a point along the beam is

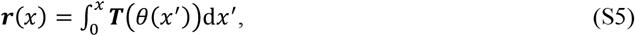

and the corresponding virtual displacement is

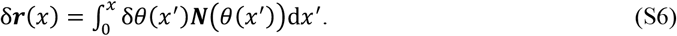

The linearized elastic potential energy of the beam and its corresponding virtual work are given by

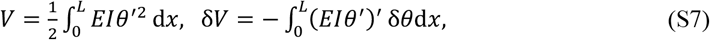

where we used partial integration and the following boundary conditions:

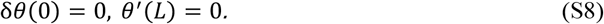

The virtual work done by the external force density ***F***(*θ*) is expressed as

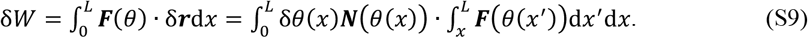

Applying the (trivial form of) d’Alemberts principle

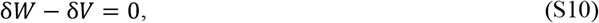

we obtain the governing equation for the equilibrium configuration:

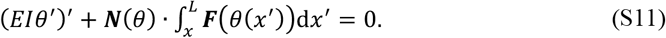

Here, the forces include contributions from multiple sources, and is defined as the sum of the acoustic excitation, streaming, frictional, and inertial forces:

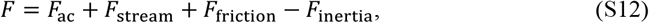

where

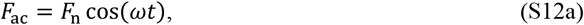

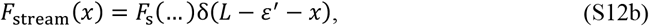

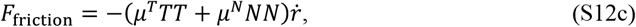

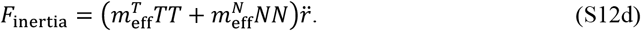

For the friction and inertial terms, we assume linear dependence on the velocity field, justified by the small-amplitude oscillation of the beam. Specifically, the beam deformation is represented as a perturbation expansion:

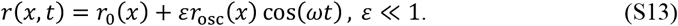

Within this perturbative framework, the streaming force *f*_*s*_ depends bilinearly on 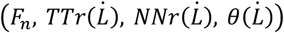 If we further assume that not only the oscillatory deformation but the total deformation is small, then the governing equation reduces to a linear ordinary differential equation for the transverse displacement *w*(*x, t*), i.e., the x-component of *r*(*x, t*).

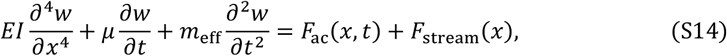

where *E* is the Young’s modulus, *I* is the area moment of inertia, *μ* is the effective viscous damping coefficient, and *m*_eff_=*ρ*_s_*A*+*C*_a_*ρ*_f_*b*^2^ is the effective mass per unit length (*3*), incorporating both the hydrogel density *ρ*_s_ and the hydrodynamic added mass via coefficient *C*_a_, with *A*=*bh* being the cross-sectional area. *F*_s_ is the streaming-induced force magnitude and *ϵ* ′ ≪ L defines the localization width. The cantilever boundary conditions are (*4*):

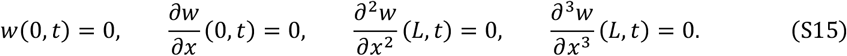

We expand the beam’s deformation into orthonormal mode shapes *ϕ*_*n*_(*x*) satisfying these boundary conditions:

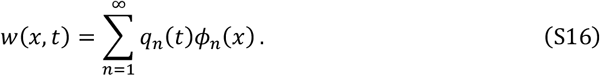

Substituting into the governing equation and applying a Galerkin projection yields a decoupled equation for each modal coordinate *q*_n_(*t*):

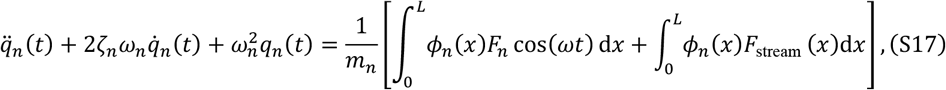

where *ω*_*n*_ is the natural frequency of mode *n*, ζ_*n*_ is the modal damping ratio, and 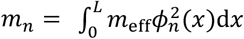 is the modal mass.

For steady-state sinusoidal excitation, the solution for each mode is:

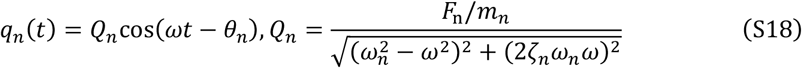

The total beam deformation is the superposition of harmonic and steady contributions:

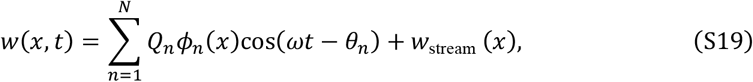

where *w*_stream_ (*x*) satisfies the static beam equation:

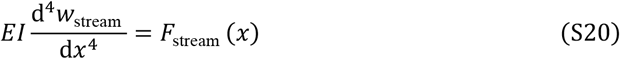

The mode shape *ϕ*_*n*_(*x*) for a cantilevered beam are expressed analytically as (*5*):

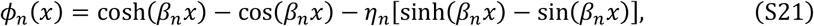

with eigenvalues *β*_*n*_ *L* determined from the characteristic frequency equation, and normalization coefficient:

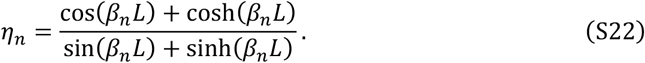

This formulation allows reconstruction of spatial and temporal deformation profiles across multiple modes.

## Supplementary Figures

**Fig. S1.**
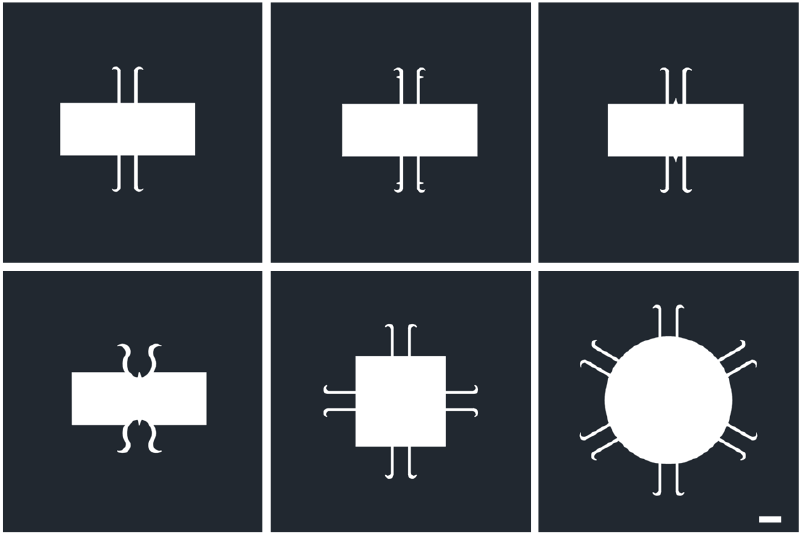
Photomasks for fabricating SonoGrippers. The fabricated SonoGrippers are scaled down by a factor of approximately16.5 relative to the dimensions on the masks designed by AutoCAD software. Scale bar, 1 mm.

**Fig. S2.**
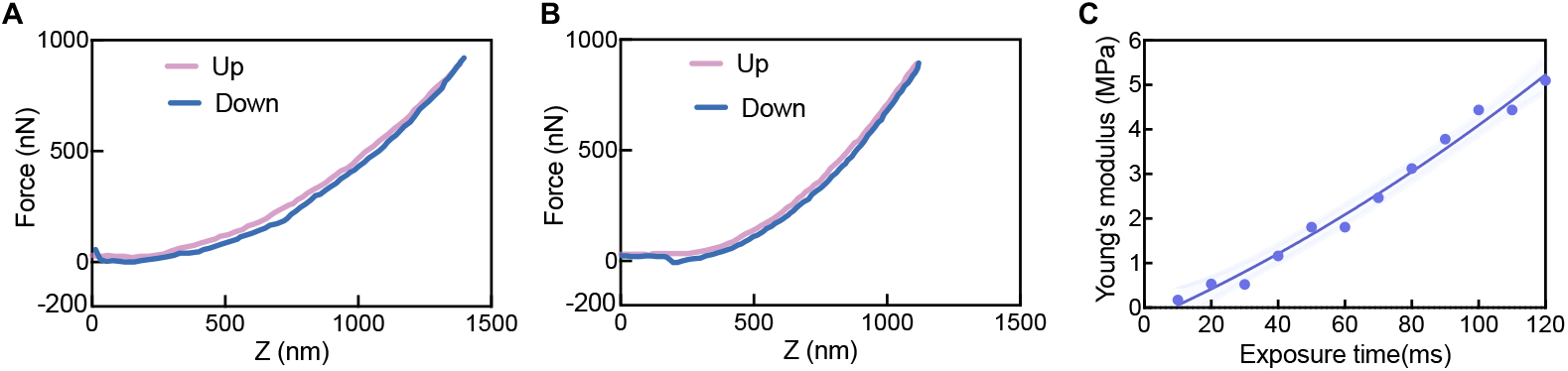
Force-displacement curves captured by AFM and the estimated Young’s modulus of SonoGrippers. Plots are force curves for SonoGrippers fabricated with UV exposure times of **(A)** 40 ms and (**B**) 50 ms. The Up and Down segments correspond to the approach and retraction phases of the AFM cantilever tip relative to the sample surface. (**C**) shows the estimated Young’s modulus of SonoGrippers fabricated with varying exposure times.

**Fig. S3.**
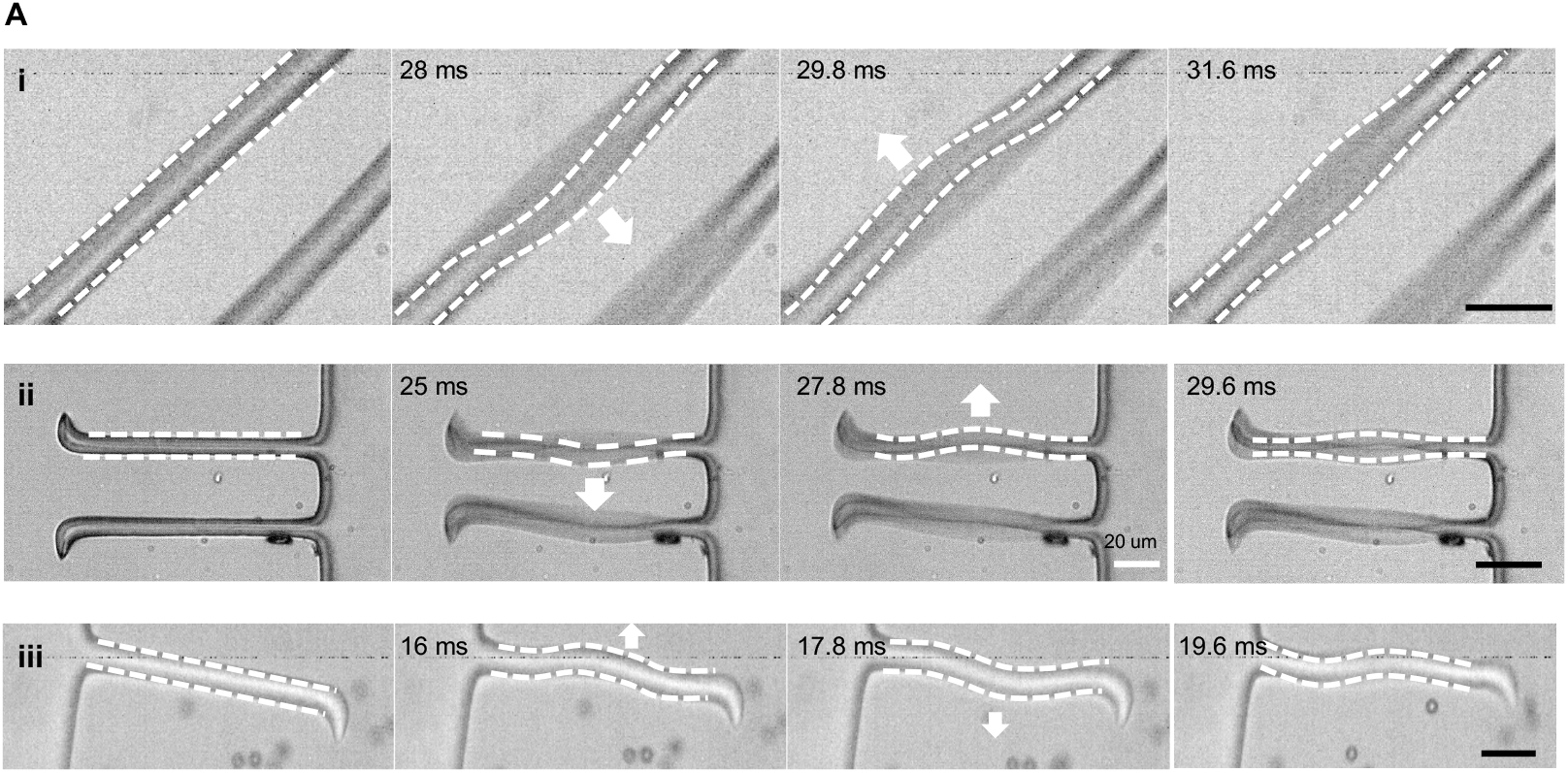
Arm deformation of Sonogrippers. Image sequences show the arm deformation of SonoGrippers with different arm lengths of i) 60, ii) 80 and iii) 100 μm. The acoustic excitation frequency was ∼5 kHz and the applied acoustic voltage ranged from 1 to 20 V_PP_. Scale bar is 20 μm.

**Fig. S4.**
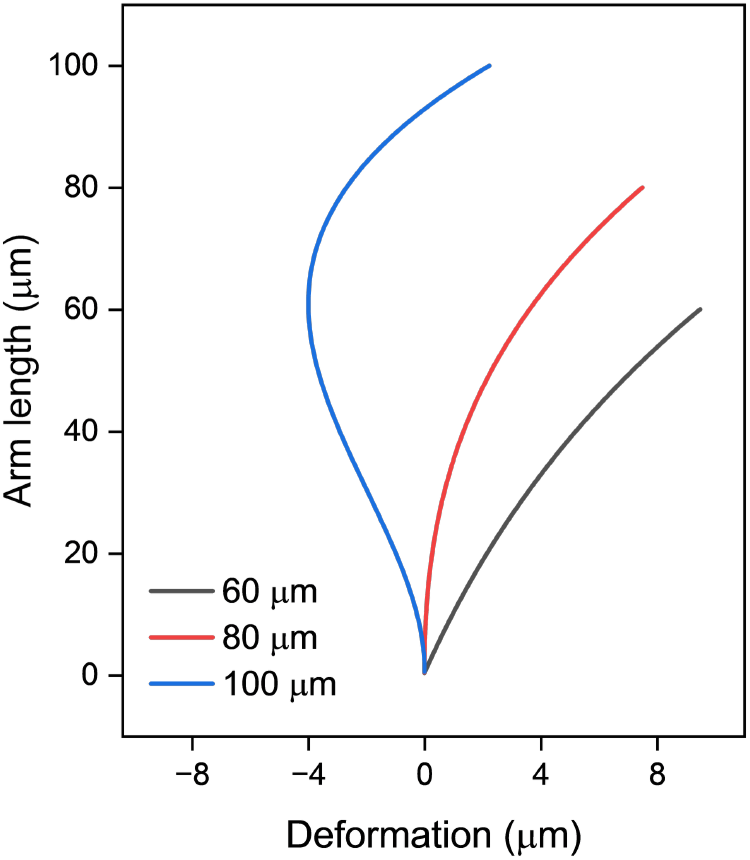
Analytical deformation profiles of cantilever arms with varying lengths. Simulated transverse deformations of soft hydrogel arms with lengths of 60, 80, and 100 μm under a 4 kHz acoustic excitation. All beams share a width of 5 μm, thickness of 20 μm, and Young’s modulus of 1 MPa. The distributed harmonic excitation has an amplitude *F*_n_=1 μN/m, and the tip-induced acoustic streaming is modeled as a localized force of magnitude *F*_stream_=10 μN/m. Longer beams exhibit more nodal structures due to increased modal participation.

**Fig. S5.**
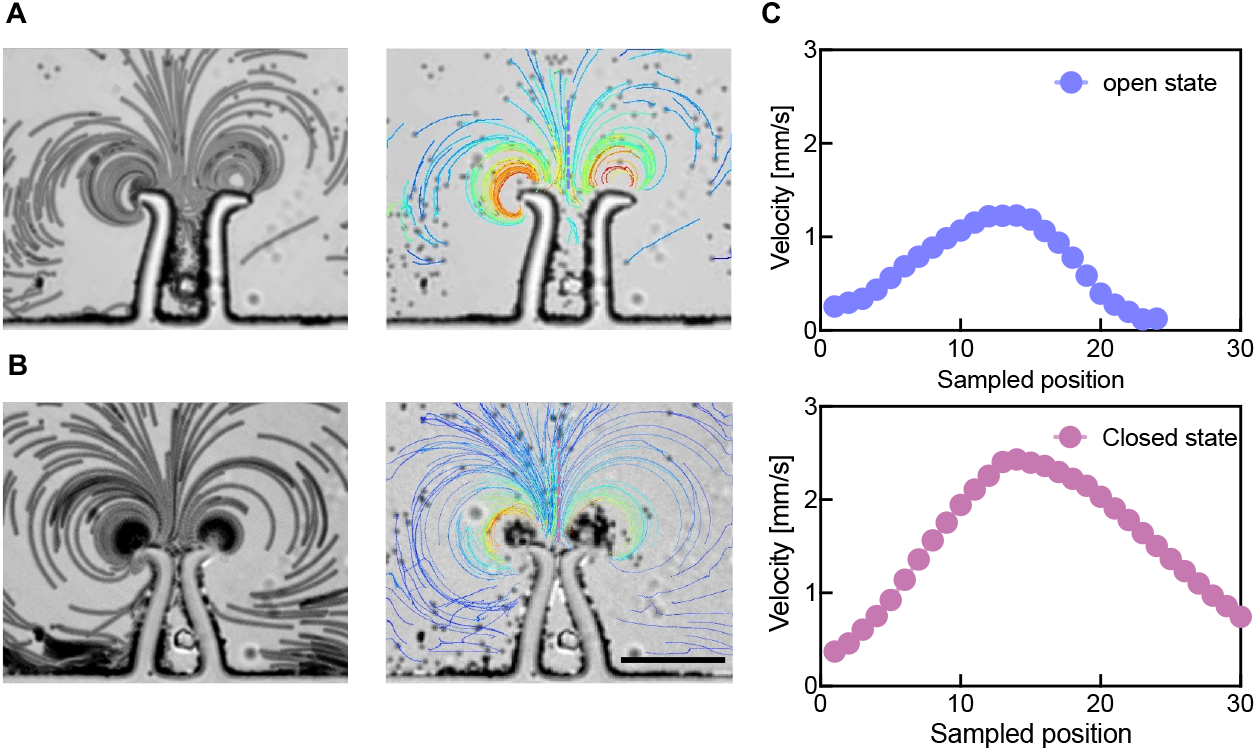
Streaming characterization of SonoGripper. Streaming profile of the SonoGripper in its (**A**) pre-closure state and (**B**) post-closure state. (**C**) Streaming velocities along the *x*-axis above SonoGrippers indicated by dash lines in (A) and (B), respectively. Scale bar is 50 μm.

**Fig. S6.**
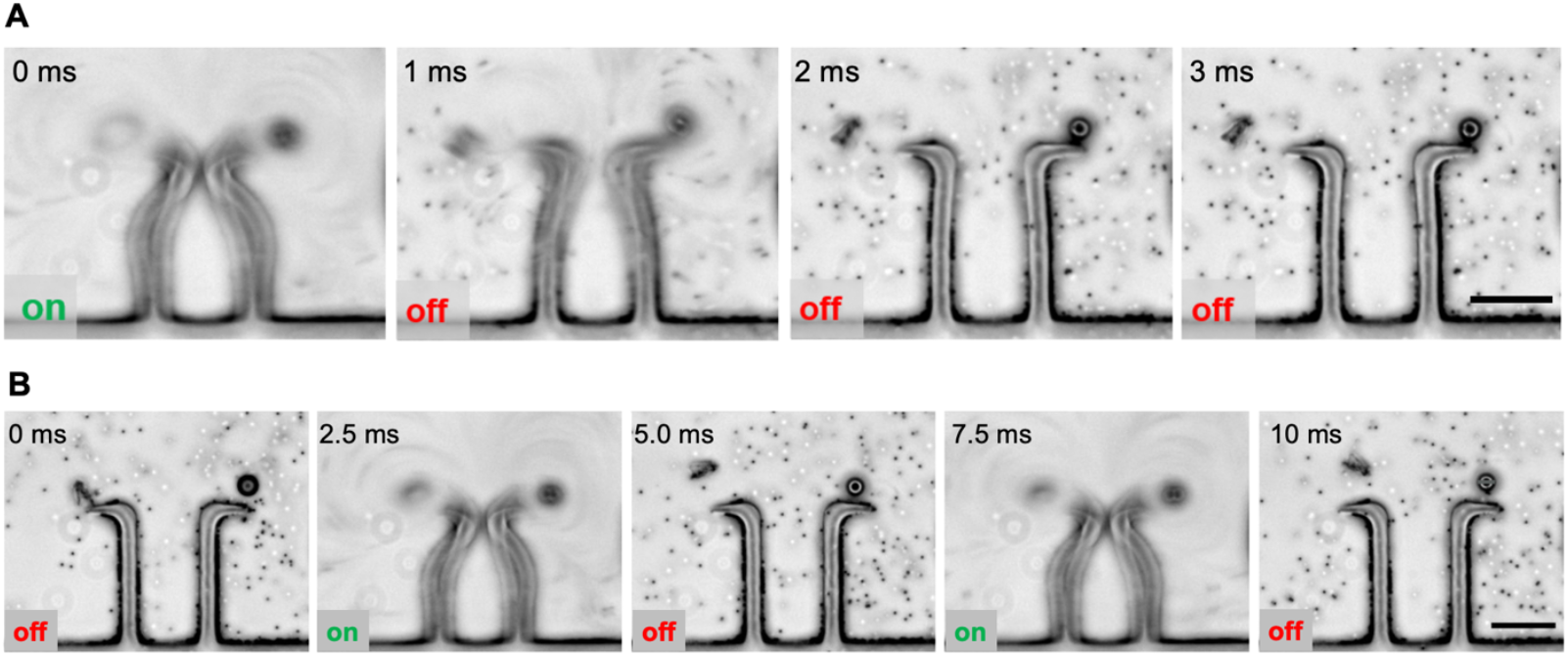
Resilient behavior and repetitive closing-opening behavior of the SonoGripper. (**A**) Resilient behavior observed upon deactivation of acoustic actuation. Time-lapse images illustrate the gripper’s recovery as the acoustic field dissipates, with its arms returning to their original equilibrium positions due to internal elastic energy storage. (**B**) Repetitive closing and opening behavior of the SonoGripper. The gripper undergoes cyclic actuation in response to the acoustic signal being turned on and off, demonstrating consistent and repeatable operation. SonoGripper was immersed in a 2 μm bead solution containing a few 15 μm beads. Scale bar is 50 μm.

**Fig. S7.**
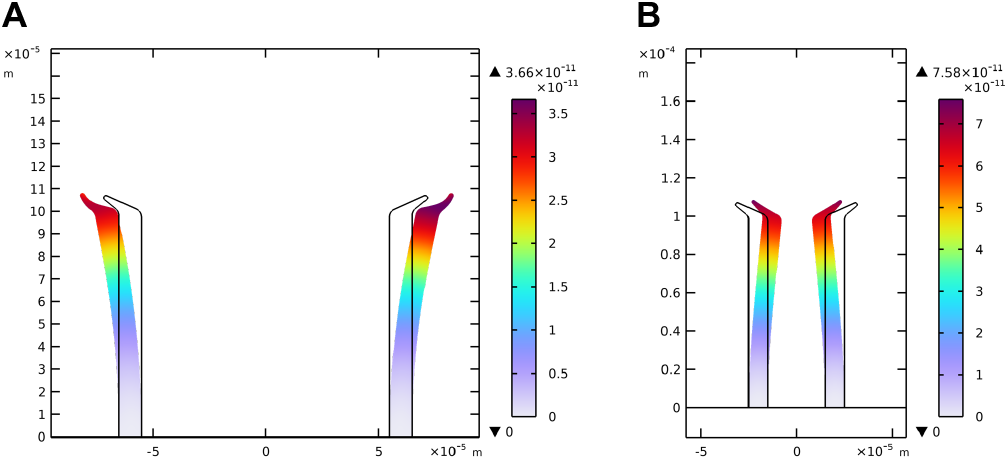
Influence of the initial arm separation distance on closure dynamics. (**A**) has an initial arm separation of 12x SonoGripper width, while (**B**) has a separation distance of 4x SonoGripper width. The color within the SonoGripper arms depicts the simulated displacement. The observed differences in simulation results demonstrate the initial arm separation distance affects the closure behavior of the SonoGripper, highlighting its tunability through geometric design modifications to meet diverse gripping requirements.

**Fig. S8.**
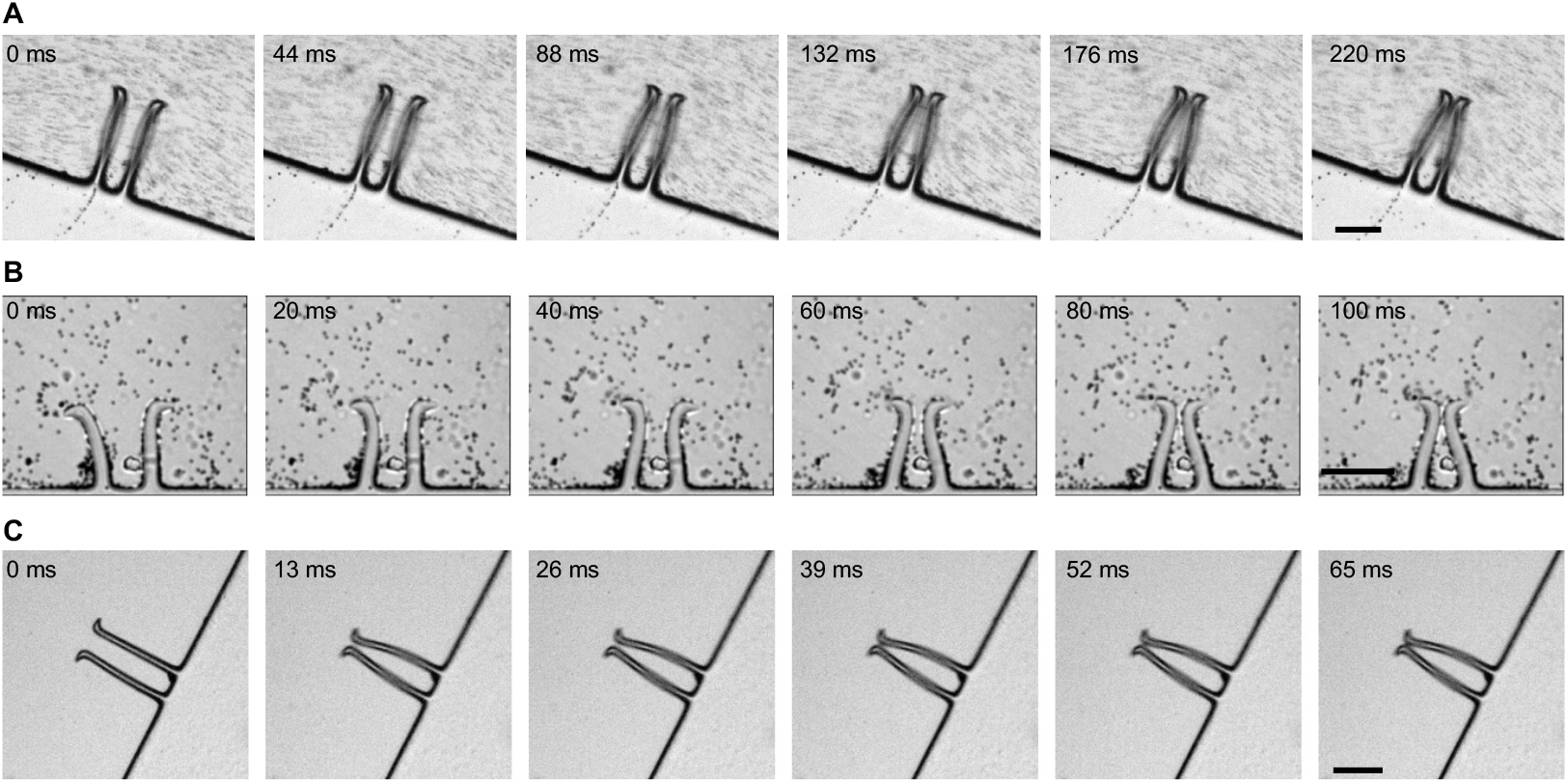
Closure process of SonoGrippers. (**A–C**) Image sequences illustrate the closure processes of different SonoGrippers with varying closure times, regulated through structural design and applied voltage control. The gripper arm lengths are 100, 60, 100 μm, while the applied voltages are 14, 16, 20 V_PP_, respectively. Scale bar is 50 μm.

**Fig. S9.**
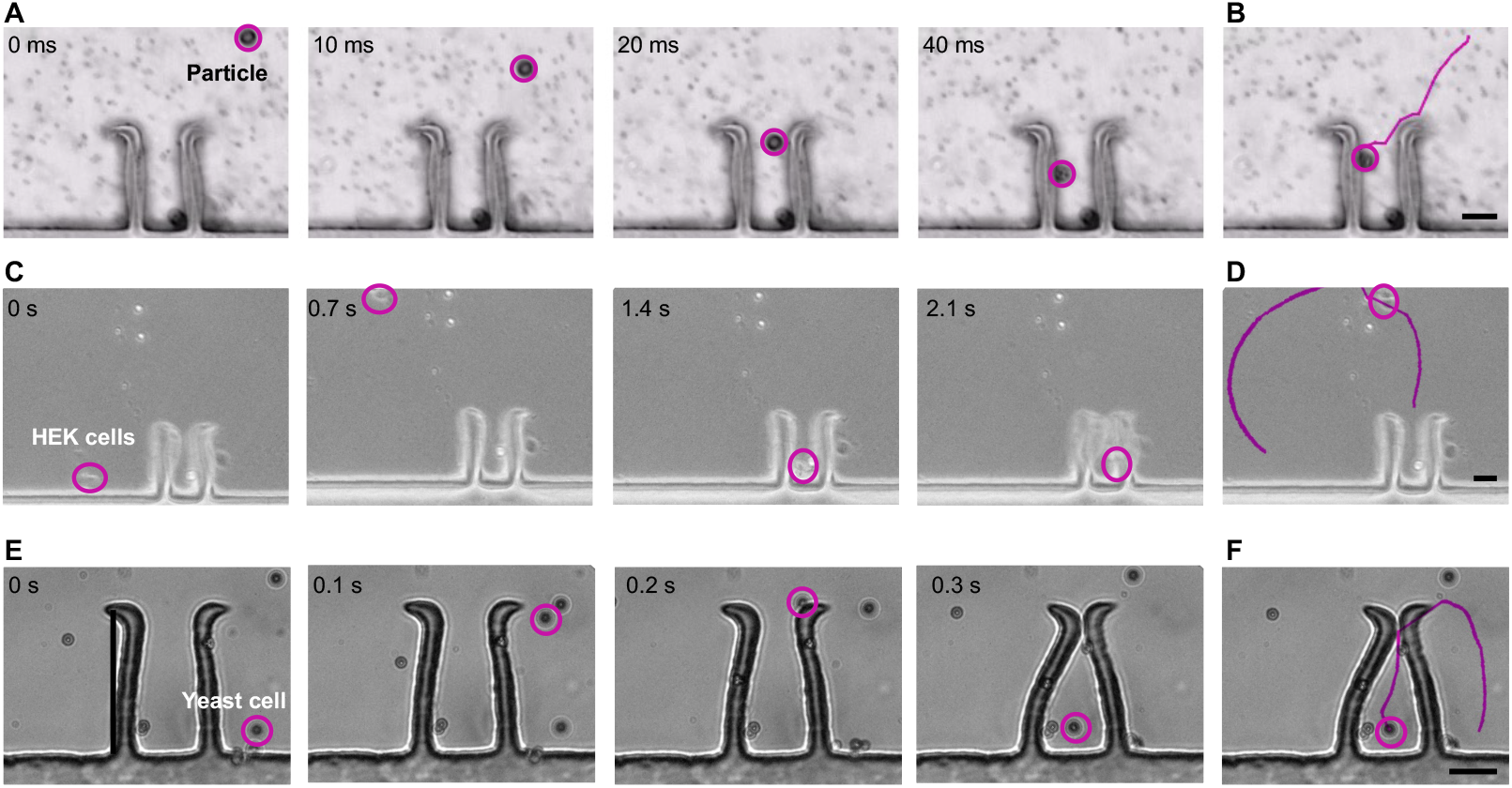
Attraction and gripping of different objects using SonoGripper. (**A**) Image sequences showing the SonoGripper capturing a 15 μm particle within 40 ms. (**B**) Tracked trajectory of the particle using the TrackMate plugin in ImageJ. (**C**) Image sequences illustrating the SonoGripper gripping HEK cells and closing its arms within 2.1 s. (**D**) Tracked trajectory of the HEK cell using the TrackMate plugin in ImageJ. (**E**) Image sequences showing the SonoGripper gripping a yeast cell within 0.3 s. (**F**) Tracked trajectory of the yeast cell using the TrackMate plugin in ImageJ. Scale bar is 20 μm.

**Fig. S10.**
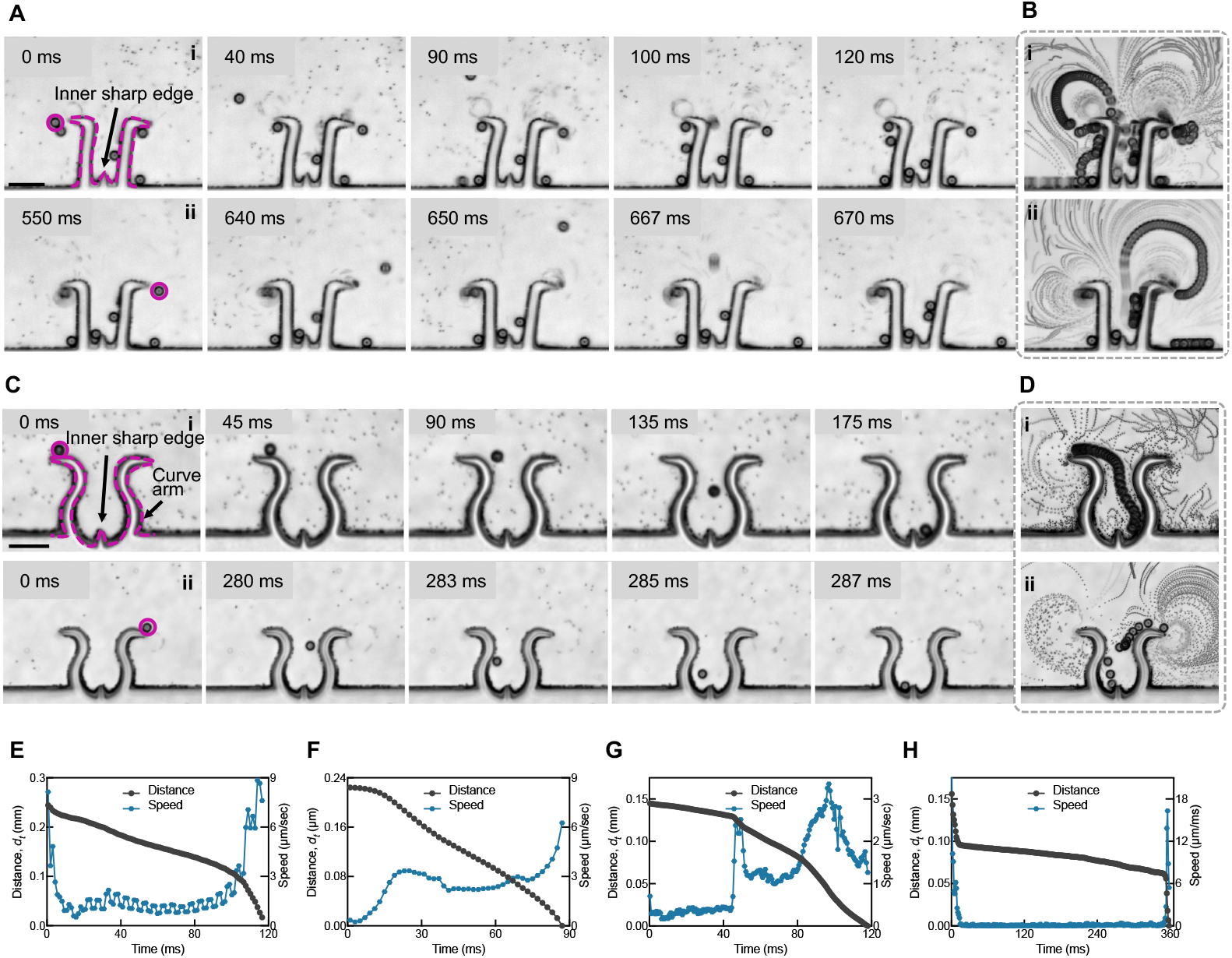
Enhanced attracting and gripping behavior of SonoGrippers with an inner sharp edge and curved arms with an inner sharp edge. (**A**) Sequential images showing a SonoGripper with an inner sharp edge attracting and gripping 15 μm particles from the left (i: 0 to 120 ms) and right (ii: 550 to 670 ms) while immersed in a 2 μm bead solution containing a few 15 μm beads and actuated at 94 kHz and 17 V_PP_. (**B**) Superimposed images of the gripping processes from the left and right, corresponding to (A). (**C**) Sequential images showing a SonoGripper with curved arms and an inner sharp edge attracting and gripping 15 μm particles from the left (iii) and right (iv). (**D**) Superimposed images of the gripping processes corresponding to (C). (**E–H**) Plots depicting the evolution of the particle distance over time and speed of the attracted particle in different trials, employing different gripper designs and approaching from the left and right directions. Scale bar is 50 μm.

**Fig. S11.**
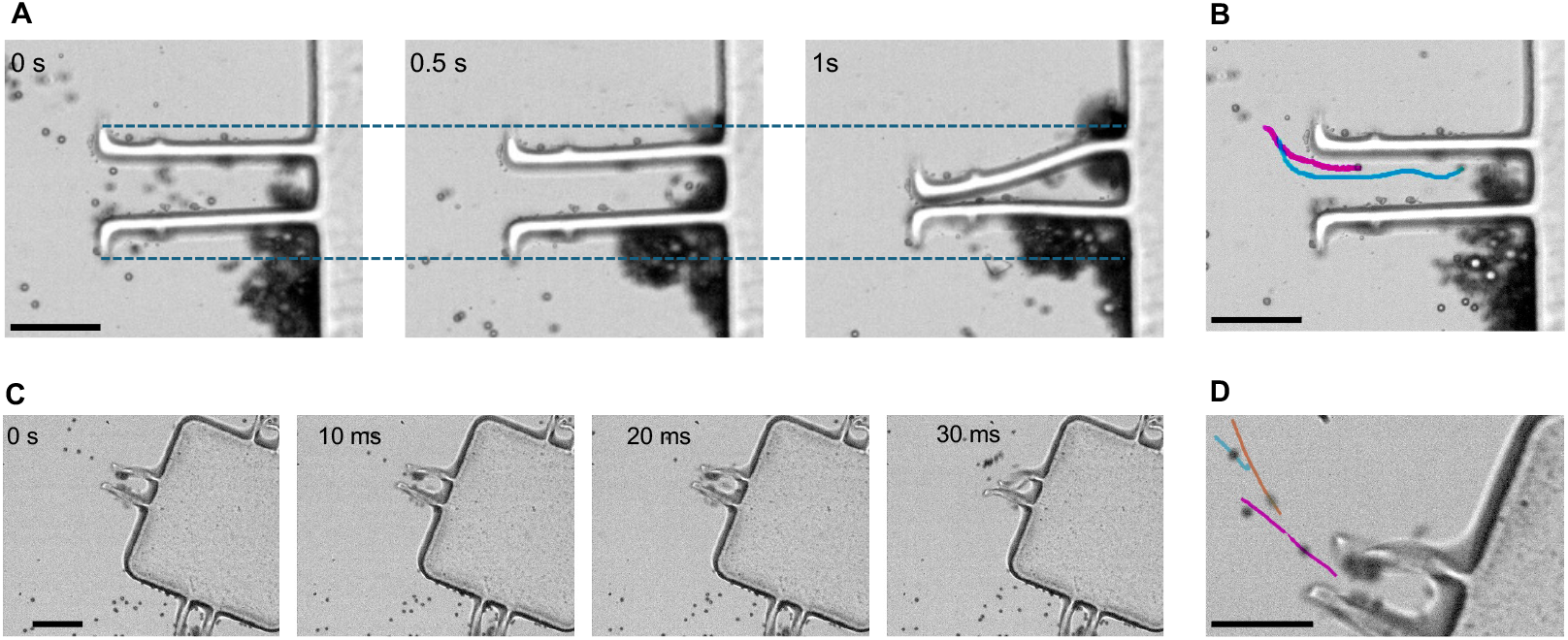
Attracting and gripping behavior of SonoGrippers with double edges and combined structures. **(A)** Sequential images showing a SonoGripper with double edges gripping two 15 μm particles and closing its arms within 1 s. **(B)** Tracked trajectories of the gripped particles using the TrackMate plugin in ImageJ. **(C)** Sequential images demonstrating the selective actuation of multi-SonoGripper array for object gripping. **(D)** Tracked trajectories of the gripped particles using the TrackMate plugin in ImageJ. Scale bar: 50 μm.

## Supplementary Tables

**Table S1.** Parameters of SonoGripper and acoustic excitation.

| Items | Parameters |
| --- | --- |
| Mixture volume ratio | 80vol%:20 vol%; 85 vol%:15 vol%; 90 vol%:10 vol% |
| UV exposure time | from 20 ms to 120 ms with an incremental step of 10 ms |
| Gripper's arm length | 50~120 $\mu\text{m}$ |
| Gripper's width | 5~20 $\mu\text{m}$ |
| Gripper's arm thickness | 20~25 $\mu\text{m}$ |
| Acoustic frequency | 5-100 kHz |
| Acoustic voltage | 1-20 V<sub>PP</sub> |

**Table S2.**
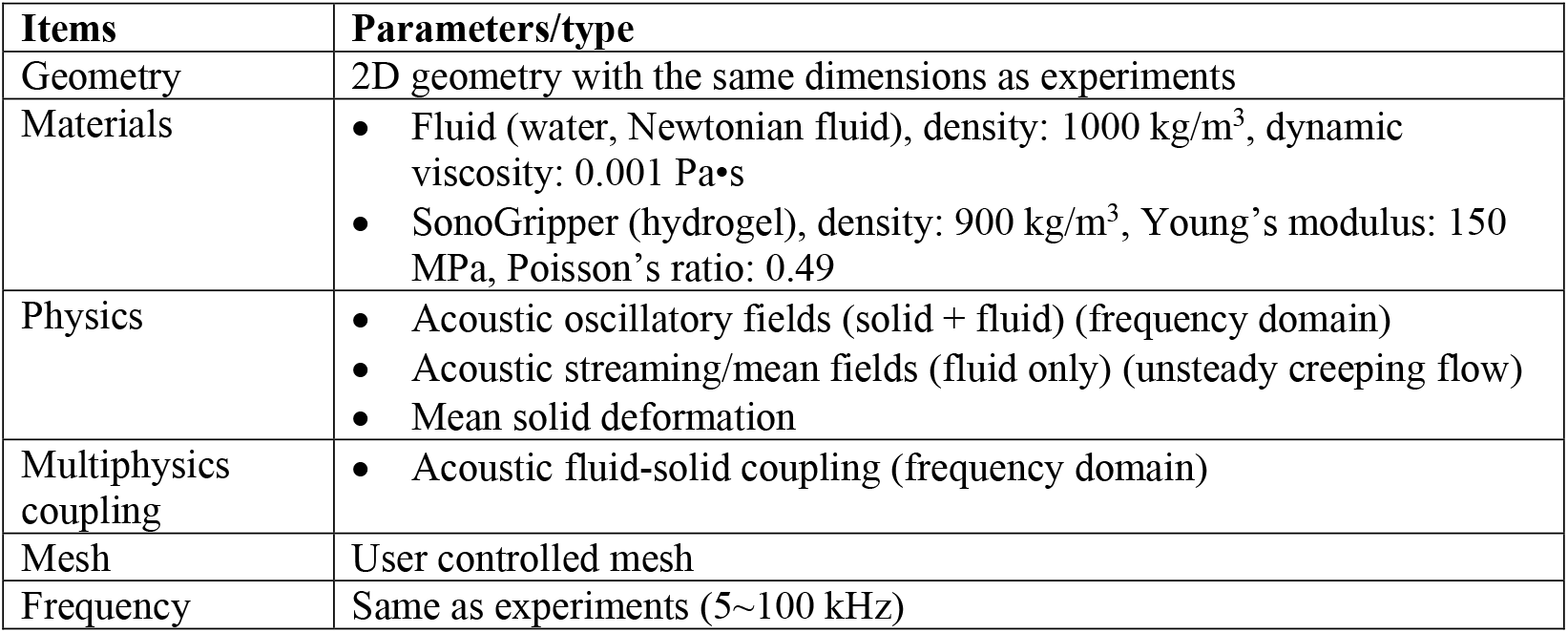
Configuration for the COMSOL simulation. Items Parameters/type.

| **Items** | **Parameters/type** |
| --- | --- |
| Geometry | 2D geometry with the same dimensions as experiments |
| Materials | • Fluid (water, Newtonian fluid), density: 1000 kg/m<sup>3</sup>, dynamic viscosity: 0.001 Pa•s<br>• SonoGripper (hydrogel), density: 900 kg/m<sup>3</sup>, Young's modulus: 150 MPa, Poisson's ratio: 0.49 |
| Physics | • Acoustic oscillatory fields (solid + fluid) (frequency domain)<br>• Acoustic streaming/mean fields (fluid only) (unsteady creeping flow)<br>• Mean solid deformation |
| Multiphysics coupling | • Acoustic fluid-solid coupling (frequency domain) |
| Mesh | User controlled mesh |
| Frequency | Same as experiments (5~100 kHz) |

## Legends for Supplementary Movies

**Movie S1: Deformation observation for SonoGrippers with varying arm lengths**. SonoGrippers were fabricated on glass substrates and activated by acoustics. An inverted microscope and a high-speed camera captured the rapid and periodic deformation behavior of SonoGrippers with arm lengths ranging from 60 μm to 100 μm. The acoustic excitation frequency was ∼5 kHz and the applied acoustic voltage ranged from 1 to 20 V_PP_. The video was recorded at 1069 and 4434 fps, and played at 30 and 10 fps.

**Movie S2: Streaming observation before and after SonoGripper’s closure**. SonoGrippers were immersed in a 2 μm tracer bead solution to visualize streaming patterns and were triggered by acoustics at 92 kHz and 16 V_PP_ to induce gradual arm closure. The transition between open and closed states was characterized by tracking tracer bead trajectories using the TrackMate plugin in ImageJ, and the resulting images were superimposed using the Z-projection function in ImageJ. The video was recorded at 4434 fps and played at 25 fps.

**Movie S3: SonoGripper closure triggered by acoustic excitation**. SonoGrippers were fabricated on glass substrates and activated by acoustics, with an inverted microscope and high-speed camera to record the closing processes of different sonogrippers. The sonogrippers have arm lengths of 105, 60, and 96 μm and were activated by acoustic frequencies of 5.1, 92, and 24.5 kHz and voltages of 14, 16, and 20 Vpp, in order. These SonoGrippers exhibited varying closure times of 220, 100, and 65 ms, respectively. The Videos are recorded at 1069 and 4434 fps, and played at 30 and 25 fps.

**Movie S4: Repetitive closing and opening of SonoGripper**. The SonoGripper undergoes cyclic closing and opening in response to the acoustic signal being turned on and off. The SonoGripper was immersed in a 2 μm bead solution containing a few 15 μm beads. The SonoGripper has arm length of 100 μm. The acoustic excitation frequence is 36.5 kHz and acoustic voltage is 20 V_PP_. The video is recorded at 1069 fps and played at 100 fps.

**Movie S5: Attracting and gripping of different objects using SonoGripper**. SonoGrippers have arm lengths of 60 μm, which are triggered under acoustic excitations with frequencies of 36.5, 6.1, and 92 kHz, respectively. In the first trial, acoustic actuation was initiated at 0 ms with an initial voltage of 10 V_PP_, which was gradually increased. Once the particle entered the gripper, the applied voltage was increased to 20 V_PP_ to induce closure. After closure, the voltage was decreased to reopen the gripper. In the trial for HEK cell gripping, the acoustic voltage was set to 5∼10 V_PP_. In the yeast cell gripping trial, the SonoGripper was actuated with a variable voltage of 1∼15 V_PP_ to facilitate attraction and gripping. The videos are recorded at 4434 fps and played at 25 fps.

**Movie S6: Attraction and gripping behavior of sonoGrippers with microstructures**. The SonoGripper with an inner sharp edge was actuated at 94 kHz with an applied voltage of 17 V_PP_, while the SonoGripper with curved arms and an inner sharp edge was actuated at 94 kHz with a voltage of 7 V_PP_. The videos were recorded at 1069 fps and played at 30 or 150 fps.

